# Models of underlying autotrophic biomass dynamics fit to daily river ecosystem productivity estimates improve understanding of ecosystem disturbance and resilience

**DOI:** 10.1101/2023.04.11.535773

**Authors:** Joanna R. Blaszczak, Charles B. Yackulic, Robert K. Shriver, Robert O. Hall

**Author notes:** Corresponding author; 1664 N. Virginia St., Mailstop 186, Reno, NV 89557. <u>Statement of authorship:</u> JRB, CBY, RKS, and ROH all contributed to model conceptualization and development. JRB wrote the first draft of the manuscript, and all authors contributed substantially to revisions. <u>Data accessibility statement:</u> All data and metadata used in this study are published and publicly available through ScienceBase from Appling *et al*. (2018b), Savoy & Harvey (2021a), and Blaszczak *et al*. (2021). All code is available at https://github.com/jrblaszczak/RiverBiomass. Author email addresses: Joanna R. Blaszczak,; Charles B. Yackulic,; Robert K. Shriver,; Robert O. Hall, Jr.

## Abstract

Directly observing autotrophic biomass at ecologically relevant frequencies is difficult in many ecosystems, hampering our ability to predict productivity through time. Since disturbances can impart distinct reductions in river productivity through time by modifying underlying standing stocks of biomass, mechanistic models fit to productivity time series can infer underlying biomass dynamics. We incorporated biomass dynamics into a river ecosystem productivity model for six rivers to identify disturbance flow thresholds and understand the resilience of primary producers. The magnitude of flood necessary to disturb biomass and thereby reduce ecosystem productivity was consistently lower than the more commonly used disturbance flow threshold of the flood magnitude necessary to mobilize river bed sediment. The estimated daily maximum percent increase in biomass (a proxy for resilience) ranged from 5% to 42% across rivers. Our latent biomass model improves understanding of disturbance thresholds and recovery patterns of autotrophic biomass within river ecosystems.

## 1 Introduction

Ecological processes vary through time in response to exogenous and endogenous factors acting over multiple time scales. Many of these factors can be directly observed and measured, such as light availability and weather conditions that can be measured using meteorological instrumentation. However, many other factors cannot be directly observed or measured either because they are not directly observable (e.g., belowground plant biomass), challenging to observe at ecologically relevant spatial and temporal scale (e.g., photosynthesis at global scales), or direct measurement would require destructive sampling (e.g., algal biomass). In cases when ecological states and processes cannot be directly observed, mathematical and statistical models are powerful tools to estimate unobserved (i.e., latent) states and processes given observable ones. For example, ecosystem ecologists use mass balance approaches to solve for unmeasured fluxes and pools given observed ones (Bormann & Likens, 1967). Animal ecologists recognize that it is challenging or impossible to directly observe the full size of a population because of imperfect detection, and thus account for detection probability to estimate population size (Williams *et al*., 2002). These approaches leverage ecological knowledge (represented in mathematical equations) and observable states and processes to infer unobservable ones, making models useful tools to improve ecological understanding not just prediction (Currie, 2019; Rastetter, 2017; Tredennick *et al*., 2021).

Advances in Bayesian methods have accelerated our ability infer unobserved processes because we can treat the unobserved states in models (i.e., latent states) as parameters to infer and account for uncertainty (Hobbs & Hooten, 2015; Auger-Méthé *et al*., 2021). A recent proliferation of carbon flux estimates enables investigation of controls on autotrophic biomass that underlies patterns in ecosystem-level gross primary production (GPP) through time. At a fundamental level, primary production depends on light availability (Farquhar *et al*., 1980; Jassby & Platt, 1976), which increases primary production over seconds to minutes by providing energy to existing autotrophic cells and over longer time periods by facilitating the accrual of autotrophic biomass. Over time scales of days to years, the relationship between light and whole ecosystem photosynthetic fluxes reflects the effects of antecedent endogenous (e.g., density dependent growth) and exogenous (e.g., disturbances) factors on current processes (i.e., ecological memory sensu (Ogle *et al*., 2015)) that change standing stocks of autotrophic organisms (i.e., biomass) (Dietze, 2017; Fisher *et al*., 2018; Moorcroft *et al*., 2001). Therefore, models of ecosystem productivity dynamics that consider both endogenous ecological feedbacks and external forcing can provide enhanced understanding of cumulative ecosystem responses to environmental change over longer time scales.

The magnitude and timing of flow and light availability explain much of the spatial variation in annual estimates of productivity among rivers (Bernhardt *et al*., 2022; Savoy & Harvey, 2021b). However, cumulative annual estimates do not reflect sub-annual variation in productivity (e.g., spring-peaks, summer-peaks (Savoy *et al*., 2019; Mejia *et al*., 2019)) that represents how underlying autotrophic populations respond to disturbance (O’Connor *et al*., 2012) and density limitations. The return interval of large disturbances that remove most autotrophic biomass is much shorter in streams than most terrestrial ecosystems (Resh *et al*., 1998; Grimm & Fisher, 1989), selecting for autotrophic species with short generation times and promoting relatively high ecosystem resilience (i.e., fast recovery) in algal biomass and primary productivity (Stanley *et al*., 2010; Grimm & Fisher, 1989; Qasem *et al*., 2019; Van Looy *et al*., 2019). Nonetheless, there are cellular limits to how quickly biomass can be produced and recovery of GPP following a disturbance lags behind the return of light availability at the stream bed. This lag creates hysteresis in the relationship between light and GPP (Fig. 1; O’Donnell & Hotchkiss (2022); Peipoch & Ensign (2022); Reisinger *et al*. (2017)). Thus, days with high light availability can yield low GPP, because of low autotrophic biomass following a flood. Biomass removal should occur during high flows when flow mobilizes sediment on the river bed (i.e., critical discharge, Wolman & Miller, 1960; Peckarsky *et al*., 2014; O’Connor *et al*., 2012), but disturbance flow thresholds based on reach-scale geomorphology are often higher than those that remove periphyton in flume studies (Hondzo & Wang, 2002). Therefore, quantifying disturbance flow thresholds that disrupt productivity based on time series of GPP may provide a more ecologically-relevant disturbance threshold estimates for rivers (Doyle *et al*., 2005) than estimates based on shear stresses necessary to mobilize bed sediments.

**Figure 1:**
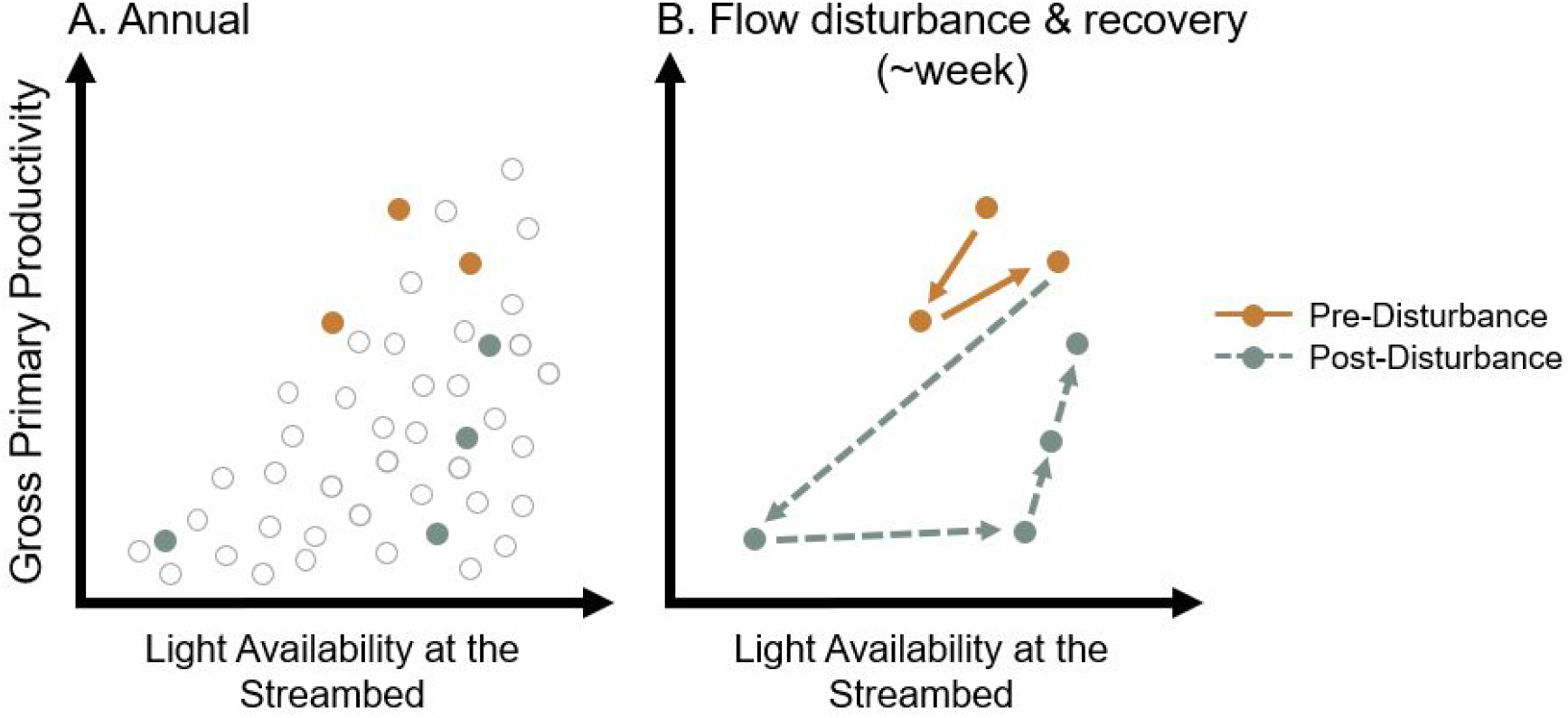
Conceptual representation of commonly observed relationships between whole ecosystem gross primary productivity (GPP) fluxes and light availability in streams and rivers on an annual time scale (Panel A) and patterns within the same data in GPP disturbance and recovery because of a high flow disturbance (Panel B). Light availability is specified to be at the streambed to account for potential light attenuation in the water column due to increased turbidity during high flows.

Early river biomass times series models with direct measures of organic matter per unit area or chlorophyll-a measurements (Uehlinger *et al*., 1996; McIntire, 1973; McIntire *et al*., 1996; Graba *et al*., 2014) enabled inferring ecological processes and model parameters at coarse resolution. However, current methods of directly quantifying river autotrophic biomass through time cannot match the spatial and temporal resolution of daily metabolism estimates in rivers because of destructive sampling techniques, complications of scaling up when biomass is patchy, and intensive labor requirements. Instead, time series models traditionally used in population ecology could advance our ability to predict GPP through time by estimating underlying variables that we cannot directly observe such as the daily variation in photosynthetically-active autotrophic biomass in a river. Here, we develop a river productivity time series model with latent biomass (LB-TS) to account for unobserved in-stream autotrophic biomass dynamics at daily temporal resolution. The biomass dynamics in the LB-TS model are in the form of a modified Ricker model (Ricker, 1954; Hobbs & Hooten, 2015), allowing for the estimation of maximum autotrophic biomass growth rates (*r_max_*) as a proxy for the resilience of primary producers to flood disturbances and the estimation of GPP disturbance thresholds in response to floods in rivers (*c*). Because increasing model complexity can result in over-fitting and poor prediction (Perretti *et al*., 2013; Ward *et al*., 2014; Tredennick *et al*., 2017; Currie, 2019), we compare the more complex LB-TS model’s predictive ability relative to a simpler standard time series model (S-TS; a first order autoregressive model). We use an inverse modeling approach to fit each model to time series of GPP from six rivers spanning different sizes and flow regimes, to answer three questions:

1. How well does a model of latent biomass dynamics including density-dependent processes recover parameter estimates and predict daily estimates of riverine GPP across an entire year relative to a simpler model?
2. How does inferred resilience (as maximal algal biomass growth rate, *r_max_*) of latent biomass vary among rivers?
3. How do disturbance flow thresholds inferred from GPP time series compare to those estimated from river hydrograph flood intervals?

## 2 Methods

### 2.1 Time series of riverine primary production

We selected six, single-station river metabolism (i.e., daily gross primary production and ecosystem respiration) time series from a broader set of 356 rivers from Appling *et al*. (2018b) (Table S1, Fig. S1). Daily metabolism estimates were generated using a state-space modeling approach (Appling *et al*., 2018a) to fit a model to sub-daily time series of dissolved oxygen (units g m*^−^*^3^), incoming light (photosynthetic photon flux density (PPFD); units *µ*mol m*^−^*^2^s*^−^*^1^) calculated from downwelling shortwave radiation from NASA’s Land Data Assimilation System (NLDAS), water temperature (*^◦^*C), depth (m), and discharge (m^3^s*^−^*^1^). The Appling *et al*. (2018a) approach used Bayesian inference to estimate the posterior probability distribution of daily gross primary production (GPP; g O_2_ m*^−^*^2^ d*^−^*^1^), daily ecosystem respiration (ER; g O_2_ m*^−^*^2^ d*^−^*^1^), and a daily standardized gas exchange rate (*K*_600_*_,d_* d*^−^*^1^). We paired all locations with attributes from the National Hydrography Database Plus version 2 (NHDv2; McKay *et al*. (2012)) using the previously published paired dataset by Blaszczak *et al*. (2021). We chose six locations across a gradient of NHDv2 Plus Strahler order river sizes (range: 2-7) that met the criteria of at least 2 consecutive years of nearly continuous daily estimates of GPP with at least 275 daily GPP estimates per year, a maximum gap of 14 d within any year, a minimal influence of canals or dams on metabolism estimates (*≤*5% of mean daily 80% oxygen turnover distances upstream of the DO sensors affected), and potential scale reduction statistics (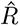*<* 1.05) for both observation error and process error parameters (Table S1-S2). Of the river locations which met this criteria (*n* = 41), we purposely chose a subset of six rivers with 2-y time series which had clear reductions in GPP following flood disturbances.

For predicting time series of GPP as described below, we compiled daily time series of daily mean discharge (*Q*) data as well as the mean and standard deviation of the posterior distributions of GPP from Appling *et al*. (2018b). We merged the GPP and Q time series with estimates of daily photosynthetically active radiation (PAR; *µ*mol m*^−^*^2^ s*^−^*^1^) at the channel surface generated by the *StreamLight* model that accounts for shading by riparian vegetation as well as clouds (Savoy & Harvey, 2021a; Savoy *et al*., 2021). Light is the primary control of photosynthesis both within rivers (Hall *et al*., 2015; Roberts *et al*., 2007) and among many rivers (Bernhardt *et al*., 2022). The effects of temperature on GPP are difficult to estimate directly because they are often mediated by nutrient availability (Welter *et al*., 2015; Huryn *et al*., 2014) or grazing (Padfield *et al*., 2017). We also do not incorporate nutrient concentrations because nutrient limitation in rivers cannot be predicted well from surface water concentrations (Keck & Lepori, 2012; Reisinger *et al*., 2016).

### 2.2 Productivity modeling

In their most general form, models describing algae growth in rivers dominated by benthic production represent the factors controlling the growth and detachment (or death) of autotrophic biomass (McIntire, 1973). We expect that autotrophic biomass growth rates slow with increasing biomass following an asymptotic relationship because of density dependence or resource limitation (McIntire, 1973; McIntire *et al*., 1996; Uehlinger *et al*., 1996). Detachment, or flow-induced loss of biomass in streams, occurs because of autogenic sloughing following senescence (Graba *et al*., 2014), loss by scour during floods (Biggs, 2000; Uehlinger *et al*., 1996), or because of consumption by grazers (Vadeboncoeur & Power, 2017). We estimated daily variation in river productivity dynamics using two state-space models fit to daily GPP time series as described below. A state-space model is a type of hierarchical modeling framework which differentiates between error introduced by stochasticity due to unquantified ecological processes (i.e., process error) versus imprecision in observations of data (i.e., observation error) (Auger-Méthé *et al*., 2021). By separating these two sources of uncertainty and modeling both a state time series which aims to represent an unknown state of nature (e.g., biomass) along with an observed time series (e.g., GPP), a state-space model can more accurately estimate uncertainty and often improves predictions (e.g., Appling *et al*. (2018a)).

#### 2.2.1 Latent biomass time series model

We developed a latent biomass time series (LB-TS) model to estimate daily GPP time series during a calendar year based on hypothesized internal dynamics of density-dependent annual growth and regrowth of photosynthetically-active autotrophic organisms (e.g., algae) following flood disturbances. The LB-TS process model estimates daily growth and disturbance of autotrophic biomass as a latent state and allows for hysteresis in the relationship between light and GPP following disturbance by defining latent biomass separately from GPP.

We chose a Ricker growth function (Ricker (1954); Hobbs & Hooten (2015); Appendix A.1) to represent the discrete time step and density-dependent growth of photosynthetically-active biomass,

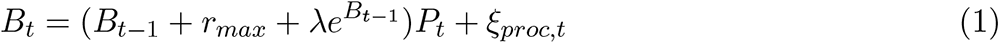

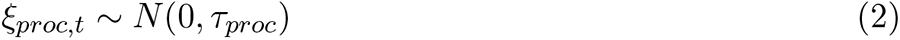

where *B_t_* is the log of latent photosynthetically-active biomass (hereafter referred to as biomass) estimated on a daily time step *t*, *r_max_* is the maximum per capita biomass growth rate, *λ* = *−r_max_/K* where *K* is the estimated biomass carrying capacity of a river, and *P_t_* is the day-to-day persistence of biomass dependent on hydrologic disturbance and detachment detailed below (Eqn. 3). The process model (Eqn. 1) represents the factors controlling biomass through time with error *ξ_proc,t_* normally distributed around a mean of 0 with standard deviation of *τ_proc_* (Eqn. 2).

We estimated the effects of flow disturbance on autotrophic biomass by modeling the day-to-day persistence (*P_t_*) of latent biomass as a function of daily mean relativized discharge (*Q_t_*). To improve parameter identifiability and model convergence, we divided each 2-y time series of daily discharge (as well as light, *L_t_*) by the annual maximum in the first year to constrain the range of relativized daily values between 0 and 1 and to maintain a consistent scaling among years when predicting the second year of data which was out-of-sample data (i.e., data not used to fit the model). We estimated daily biomass persistence (*P_t_*) using a complementary log-log link function,

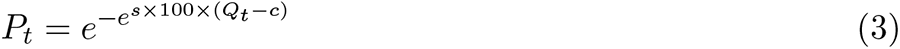

where *P_t_* is the persistence of log latent biomass *B_t_* (see Eqn. 1) from day-to-day which is a proportion of the previous day, *s* is a parameter characterizing the sensitivity of the persistence transition scaled by 100 to improve parameter identifiability, *Q_t_*is the daily mean discharge relative to the annual maximum discharge as described above, and *c* is the flow disturbance threshold parameter which approximates the *Q_t_* threshold at which latent biomass is disturbed. The log-log link function (Eqn. 3) constrains values between 0 and 1, where 0 is no persistence and 1 is complete persistence. *P_t_* has an asymptotic relationship with zero which captures the persistence of residual biomass remaining on rocks and other substrata in rivers following a flow disturbance (i.e., refugia) that allows for regrowth within a river reach (Uehlinger *et al*., 1996; Van Looy *et al*., 2019).

The observation model fit to the daily GPP data was,

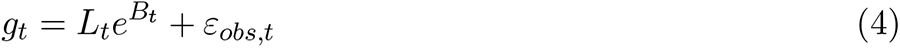

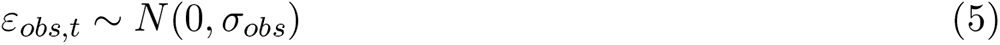

which is to say that GPP on a particular day is the product of biomass and light, where *g_t_*on a daily time step *t* is the daily mean of the posterior probability distribution (i.e., the distribution of parameter estimates given the data and our prior knowledge; hereafter referred to as posterior) of previously modeled daily GPP time series (g O_2_ m*^−^*^2^ d*^−^*^1^), *L_t_* is relativized light, and *B_t_* is the log of biomass as described in Equation 1. The observation model (Eqn. 4) has error *ε_obs,t_* normally distributed around a mean of 0 with standard deviation of *σ_obs_* (Eqn. 5). The *σ_obs_* prior (i.e., the assumed probability distribution of the parameter given our existing or lack of existing knowledge about a parameter in the model) was a normal distribution truncated at zero with a mean and standard deviation corresponding to the mean and standard deviation of the Appling *et al*. (2018b) daily GPP posterior. All other fitted parameters were assigned weakly informative priors to limit the posteriors to realistic parameter spaces (see Supporting Information Table S3 for prior assignments). We defined the initial value of *B_t_* as normally distributed around the log of the first day of GPP data for a given year divided by *L_t_* with a *σ* of 1. We did not incorporate a light use efficiency coefficient (*α*) which is often used in productivity-biomass relationships because the latent state of biomass would inherently covary with *α*. Therefore, we only retain latent biomass with the assumption that we are only measuring contributions of photosynthetically-active biomass that produce or consume oxygen measured by in-stream sensors. We were able to recover all parameter estimates when we simulated data with known parameters (Fig. S2-S3).

#### 2.2.2 Standard time series model

Introducing increasing complexity into models can often decrease model performance and result in worse predictions (Ward *et al*., 2014). Therefore, we compared the predictions from the LB-TS model to those from a simpler standard time series (S-TS) model of GPP with light and flow similar to Roley *et al*. (2014) and Hall *et al*. (2015) as a benchmark. Such a low-dimensional model does not allow for hysteresis due to biomass recovery and takes the form,

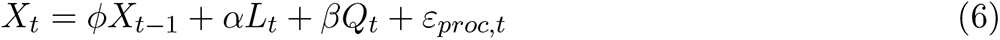

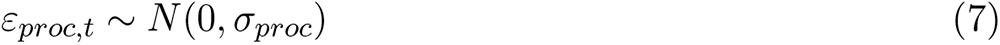

where *X_t_* on a daily time step *t* is the predicted GPP (g O_2_ m*^−^*^2^ d*^−^*^1^) on a log scale to ensure all values are positive, *φ* is an autoregressive parameter multiplied by the log of the previous day’s GPP value to represent lagged effects, *α* is a positive light use efficiency parameter of log GPP growth per unit relativized light (*L_t_*), and *β* is a negative loss parameter of log GPP loss per unit relativized discharge (*Q_t_*). The process model (Eqn. 6) represents the factors controlling GPP through time with error *ε_proc,t_* normally distributed around a mean of 0 with standard deviation of *σ_proc_* (Eqn. 7). The observation model fit to the data was,

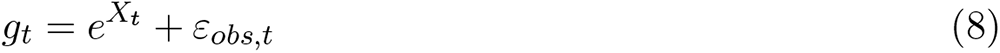

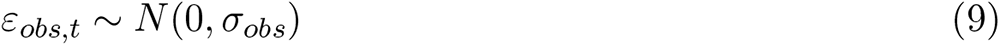

where *g_t_* is the daily mean of the posterior of previously modeled daily GPP as described in Equation 4 and the observation model (Eqn. 8) has the same error distribution for *ε_obs,t_* and assigned priors as in Equation 5. We defined the initial value of *X_t_* as normally distributed around the log of the first day of GPP data for a given year with a *σ* of 0.1. All other fitted parameters were assigned weakly informative priors to limit the s to realistic parameter spaces (see Supporting Information Table S3). As with the LB-TS model, we could recover all parameter estimates when we simulated data with known parameters (Fig. S4-S5).

### 2.3 Model implementation, diagnostics, and comparison

We fit both state-space model variants using Bayesian inference via Hamiltonian Monte Carlo in Stan (Carpenter *et al*., 2017) and the rstan interface (Stan Development Team, 2020) in R (R Core Team, 2022) (Fig. S6 – S7). We ran four chains for 5000 iterations with a 2500 iteration warm-up, a maximum tree depth of 12, and a target average acceptance probability of 0.95. First, we fit both the LB-TS and S-TS models to the first year of each 2-y time series from 6 different rivers (Table S1). We evaluated parameter convergence using trace plots and the potential scale reduction statistic (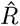) of parameters. All parameter estimates had a 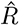*<* 1.05 (Table S4-S5).

We evaluated models based on their predictive accuracy using out-of-sample prediction tests (Hooten & Hobbs, 2015; Gelman *et al*., 2014) in which we predicted the second year of the GPP time series for each site using the posterior of the parameters from each model fit to the first year of data. We calculated the normalized root mean squared error (NRMSE) of the model predictions by dividing the root mean squared error (RMSE) of the predictions by the range in the data. We also calculated the coverage of the model predictions as a percentage of observed data points that fell between the 2.5% and 97.5% intervals of the model predictions. Finally, we compared LB-TS model parameter estimates and out-of-sample predictions for one site (South Branch Potomac River, WV) when the model was fit to a shorter one year of “training” data (as was done for all other locations) versus four years of data. We compared out-of-sample GPP predictions generated by the LB-TS and S-TS models by calculating the evolving information state of each model over the predicted year to estimate how support for either model changed during the time series (Nichols *et al*. (2019, 2021); Appendix A.2).

We compared disturbance flow thresholds inferred from the persistence parameter *P_t_* in Equation 3 to the flow threshold for incipient sediment motion on a river bed that can disturb benthic biomass. Infrequent, high magnitude flows can cause the most severe alteration of river bed morphology; however, frequent storm flows of a moderate magnitude (i.e., bankfull floods that occur with *∼*1-2 y recurrence intervals) are commonly what mobilizes the most sediment over time (Wolman & Miller, 1960; Buffington & Montgomery, 1997; Mao & Surian, 2010). The disturbance flow threshold of latent biomass as identified by the *c* parameter in Equation 3 is not meant to be equivalent to geomorphic effective discharge; instead, *c* represents the minimal magnitude of flow necessary to reduce latent biomass. However, the comparison of *c* estimates to hydrologic estimates of effective discharge are still useful for understanding how flow magnitudes that cause the most movement of bed sediments compare to the minimum necessary to reduce GPP a detectable amount. Therefore, we back calculated the discharge for each river corresponding to *c* (*Q_c_*), and we adopted two simplifying assumptions from Wolman & Miller (1960) that have mixed support in the literature (Torizzo & Pitlick, 2004; Lisenby *et al*., 2018) but allowed us to estimate effective discharge despite a lack of widespread geomorphic information for rivers at a continental scale (e.g., grain size distributions). First, we assumed that the effective discharge and the bankfull discharge of a river are approximately equal (Pfeiffer *et al*., 2017; Phillips & Jerolmack, 2016). Second, we assumed that bankfull discharge is represented by flows of recurrence intervals of 2 y. We downloaded USGS mean daily discharge data for each site from January 1, 1970 to December 31, 2020, fit a linear relationship between the log of the maximum annual discharge and the exceedance probability (Appendix A.3) using maximum likelihood estimation, and then used the relationship to estimate the magnitude of the 2-y flood recurrence interval discharge (Hornberger *et al*., 1998). We also examined how *Q_c_* varied among rivers across stream order and contributing watershed area.

Finally, we estimated the resilience of biomass within a site to hydrologic disturbance as the median value of the low-density (i.e., maximum) growth rate *r_max_* parameter from the LB-TS model. We compared the estimated *r_max_* for each site across contributing watershed area, stream order, mean incoming light at the stream surface, and mean water temperature. We calculated the daily maximum percent increase in biomass as (*e^r^^max^ −* 1)*×* 100 and the maximum doubling time of biomass as ln(2)/*r_max_*. All calculations were completed in R version 4.0.2 (R Core Team, 2022).

## 3 Results

### 3.1 Annual estimates of river productivity and latent biomass dynamics

The latent biomass time series (LB-TS) model adequately fit daily gross primary productivity (GPP) time series data based on its predictive ability of a subsequent year of data. The estimated daily variation in underlying autotrophic biomass varied with discharge (Fig. 2B, Fig. S8) while predicting most daily variation in GPP for each river. The LB-TS model out-of-sample predictions using the full posterior distributions from the model fit to the previous year (e.g., Fig. 2A,B), predicted most of the variation in daily GPP estimates for the following year across all sites (Fig. 2C, Fig. S9). The LB-TS model out-of-sample predictions tended to have the greatest bias in the spring and summer by under-predicting peak GPP (Fig. S9). A longer training data set to inform model estimates in the South Branch Potomac River, WV site provided more constrained parameter estimates for all parameters except *c*. This constraint improved prediction of low-GPP dynamics, yet further under predicted of spring peaks in GPP, likely because additional training years contained somewhat lower peak GPP than the out-of-sample year (Fig. S10).

**Figure 2:**
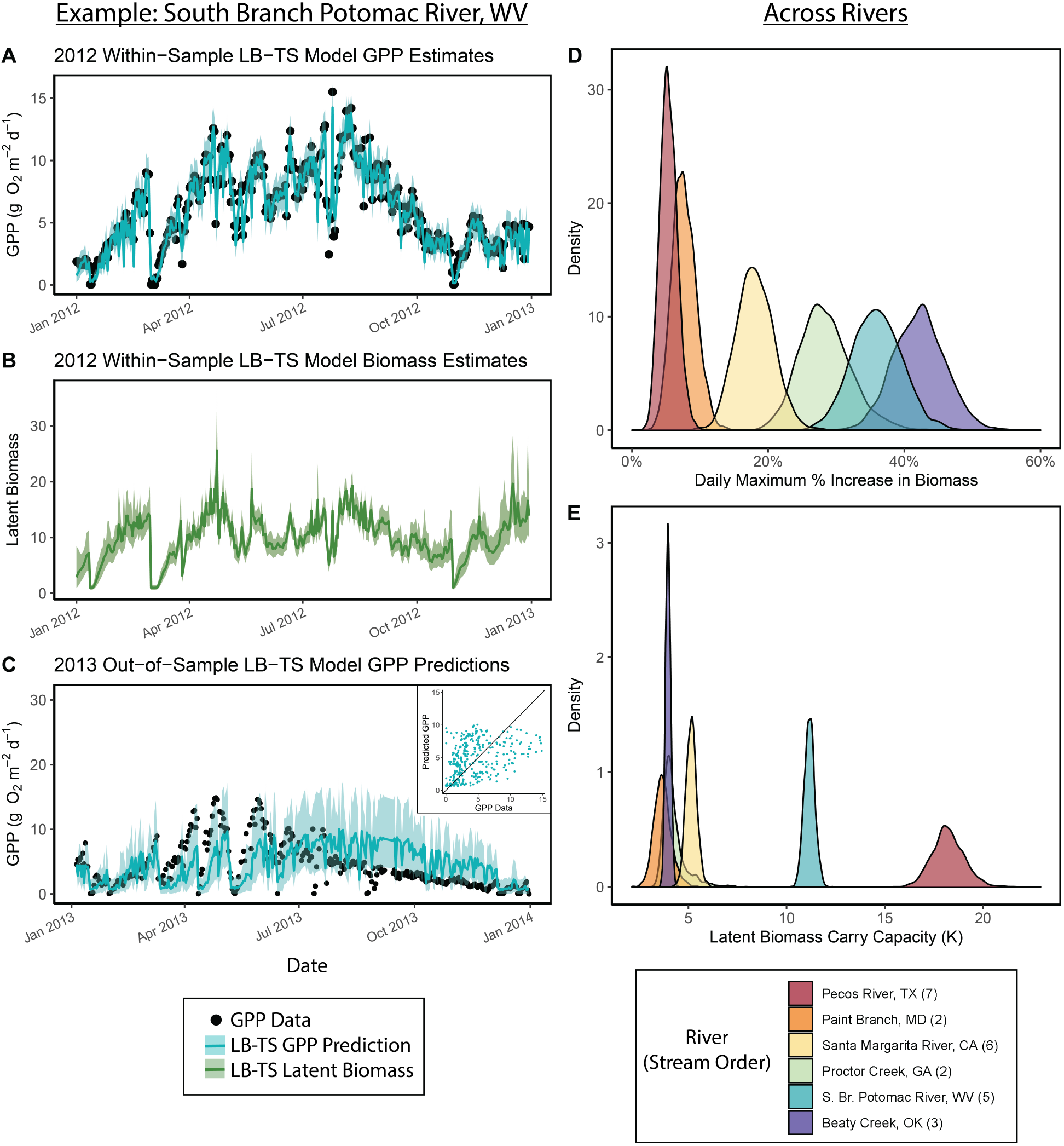
LB-TS model parameter estimates and predictions for an example river (South Branch Potomac, WV; Panels A-C) and across all six rivers (Panels D & E). Lines in Panels A & B with ribbons represent the median and uncertainty intervals (2.5% – 97.5%) of daily GPP and latent biomass LB-TS model estimates, respectively. Black dots are previously estimated GPP data. Panel C shows the median and uncertainty intervals of the LB-TS model predictions for the out-of-sample year (2013) generated using the full within-sample (2012) posterior distributions and the inset plot shows the comparison between GPP data and predicted GPP. The posterior probability distributions for the daily maximum percent increase in biomass ((*e^r^^max^ −* 1)*×* 100; Panel D) and the carrying capacity (*K*; Panel E) across all rivers colored by median within-sample *r_max_* estimates.

When compared to a simpler model (S-TS), the more complex LB-TS model had similar or improved model performance. Coverage by out-of-sample predictions was similar or better (S-TS mean coverage: 82% (range: 64-97%); LB-TS mean coverage: 87% (range: 70-94%); Fig. S11, Table S6). In addition, the LB-TS model had similar or improved mean out-of-sample GPP prediction accuracy relative to the S-TS model (LB-TS mean NRMSE: 0.21 (range: 0.15-0.29); S-TS mean NRMSE: 0.24 (range: 0.19-0.31)). Cumulative model support for the more complex LB-TS model out-of-sample predictions was strongest in Paint Branch, MD and Santa Margarita River, CA (inset plots in Fig. S11), while model support in the remaining rivers alternated between the S-TS model and LB-TS model or was approximately equal by the end of the year.

Latent biomass tracked variation in flow in rivers with build ups between storms and removal of biomass immediately following storms. Daily variation in within-sample biomass (*B_t_*) co-varied tightly with modeled GPP through time (Fig. S12). While daily *B_t_* estimates were dynamic through time when fit to GPP time series (e.g., Fig. S7), daily out-of-sample *B_t_* predictions plateaued in between disturbance events at the estimated carrying capacity (*K*) for each river (Fig. S13). The magnitude of *K* estimates (range among rivers: 3.6-18.2) increased with the median observed GPP (*r* (4) = 0.99, *p <* 0.001) reflecting the representation of *K* as a metric of the capacity of the river to sustain a particular amount of photosynthetically-active autotrophic biomass.

### 3.2 Resilience of river productivity

Rivers varied strongly in their resilience to floods. Variation in the resilience of river productivity to flow disturbances as estimated by the maximum growth rate of latent biomass (*r_max_*) was more distinct among rivers than the carrying capacity (Fig. 2D,E). Median *r_max_* estimates ranged between 0.05 (5% daily maximum increase in biomass) in the Pecos River, TX to 0.35 (42% daily maximum increase in biomass) Beaty Creek, OR (Table S4; Fig. 2D), equivalent to a range of estimated biomass doubling times of 2 to 14 days. Individual rivers had different patterns in *r_max_* and *K* (Fig. 2D,E). The largest river included in this study (Pecos River, TX) had the highest *K* while the lowest *r_max_* (slowest recovery time). A mid-sized stream (Beaty Creek, OK) had a nearly inverse pattern, with the highest *r_max_* and one of the lowest *K* estimates. And yet, one of the smallest and shadiest rivers (Paint Branch, MD) had both relatively low *r_max_* and *K* estimates. However, when investigated across our limited sample size of six sites, we did not observe clear relationships between median *r_max_* and watershed area, stream order, mean annual PAR at the stream surface, nor mean annual stream water temperature (Fig. S14).

### 3.3 Disturbance flow thresholds and post-disturbance hysteresis of river productivity

The estimated magnitude of flow necessary to remove biomass was always lower than the 2-y flood. For the first year in each site, the median estimates of critical disturbance flow thresholds (*Q_c_*) fell within or were slightly above the range of observed discharge and all were lower than the magnitude of the 2-y recurrence interval flood (*Q*_2_*_yr_*) (Fig. 3). Median *Q_c_* estimates ranged from 14% to 101% of the maximum discharge observed within the time series used to fit the model (*Q_max_*), and 3-72% of the *Q*_2_*_yr_* (Fig. S15, Fig. S16). There was no relation between the relative magnitude of *Q_c_* and stream order or the size of the contributing watershed (Fig. S17). The sensitivity (*s*) parameter in the LB-TS model represents the sharpness of the transition from nearly complete day-to-day persistence (*P_t_*) of latent biomass to the nearly complete removal of biomass following a flood disturbance. *s* estimates were similar across sites (range: 1.26 – 1.74; median: 1.5; Fig. 3) with sharp transitions between the two extremes (almost all to none) of day-to-day biomass persistence past the *Q_c_* threshold.

**Figure 3:**
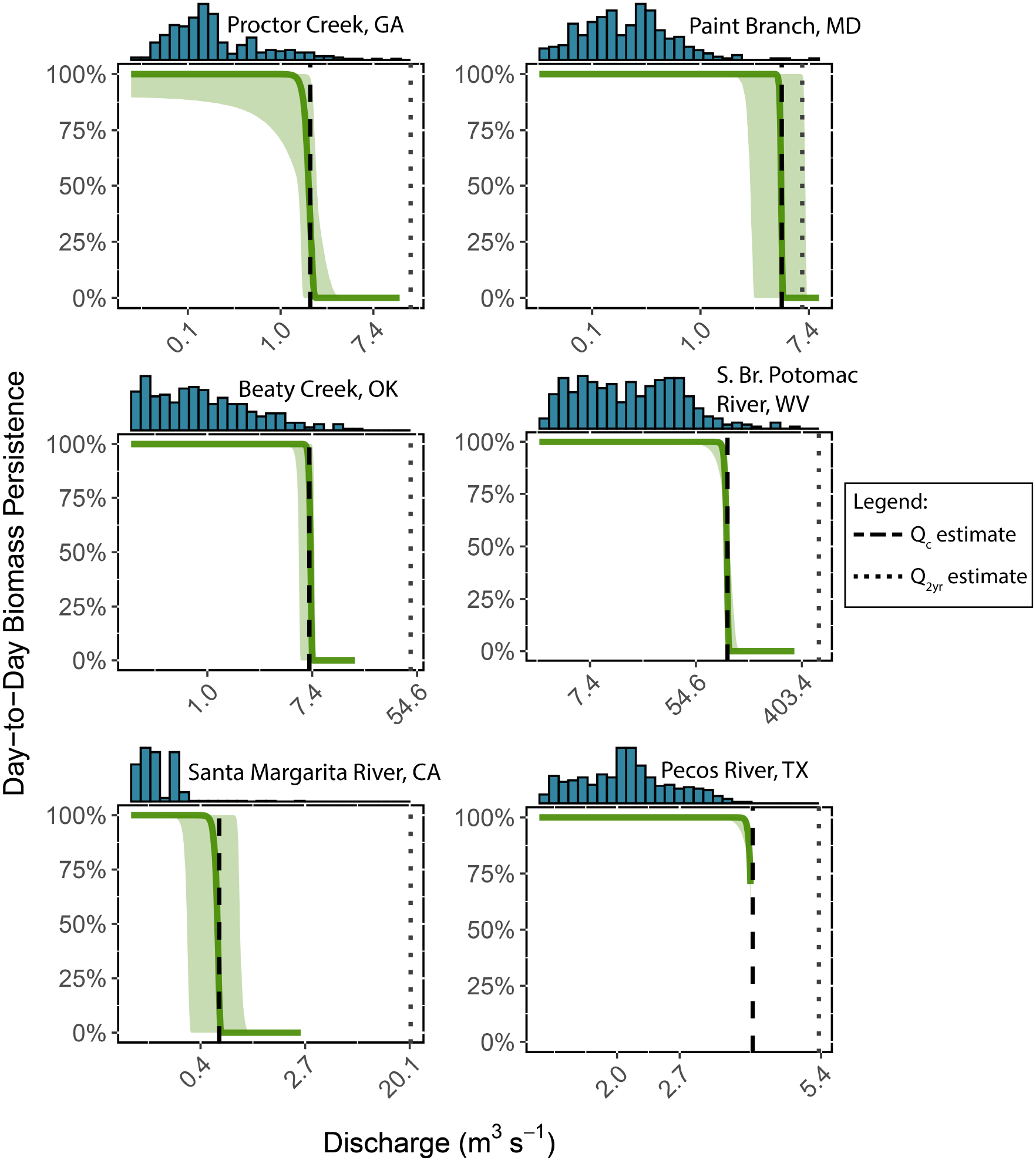
Estimated daily persistence of latent biomass *B_t_* as a percentage across the range of observed daily discharge (m^3^ s*^−^*^1^) in the within-sample year for each of the six river time series. A histogram of the observed daily discharge is shown at the top of each plot corresponding to the x-axis. Green lines with ribbons represent the median and uncertainty intervals (2.5% – 97.5%) of the simulated *P_t_* across the range of observed mean daily discharge within each site calculated using Equation 3 from the full posterior distributions of the discharge threshold for latent biomass disturbance *c* and the sensitivity of the disturbance threshold *s*. Green lines are truncated at the upper limit of observed flow. Complete day-to-day persistence approximates 1 (100%), while near complete removal approximates 0 (0%). The median critical flow discharge (median *c* estimates converted to the corresponding daily discharge (m^3^ s*^−^*^1^) value) for each site is shown as a vertical dashed line and the estimated 2 y recurrence interval flood is shown as a vertical dotted line.

Out-of-sample predictions of latent biomass following flow disturbances exhibited hysteresis patterns between latent biomass and light availability. We highlight one example during May 2013 in the South Branch of the Potomac River, WV. In the week prior to high flows on 8 May 2013, daily variation in GPP was primarily controlled by variation in light (Fig. 4A, B). Following the disturbance, variation in GPP was decoupled from light and instead the recovery of latent biomass was the dominant signal until GPP returned to a similar magnitude as before the disturbance (Fig. 4C, D, E).

**Figure 4:**
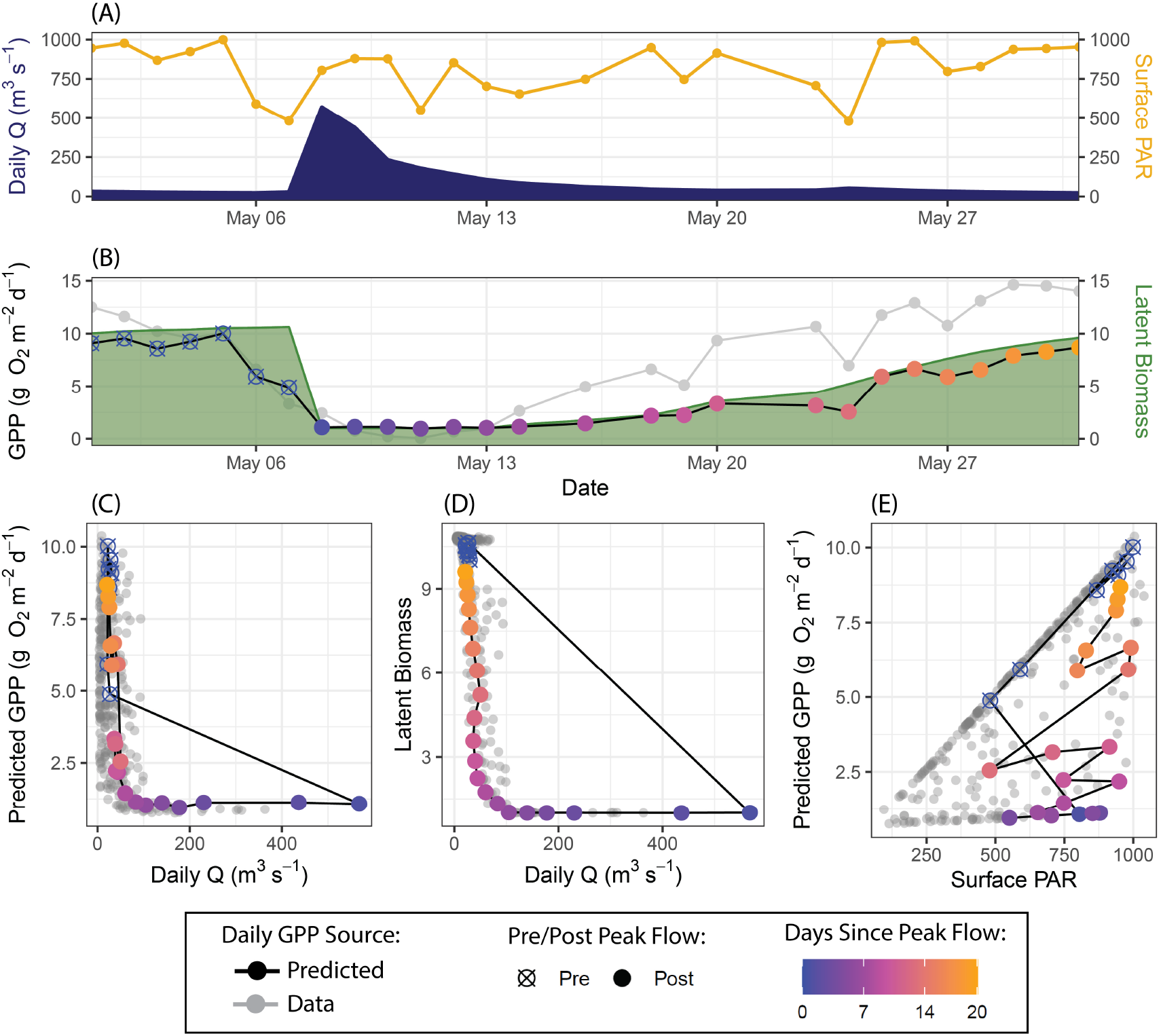
Example of hysteresis patterns between light availability and daily gross primary productivity (GPP; g O_2_ m*^−^*^2^ d*^−^*^1^) in the South Branch of the Potomac River, WV from 1-31 May, 2013 following high flows on 8 May. (A) Surface photosynthetically active radiation (PAR; *µ* mol m*^−^*^2^ s*^−^*^1^) in yellow and daily discharge (Q; m^3^ s*^−^*^1^) in blue. (B) Time series of original GPP estimates (grey), predicted GPP estimates (black lines with point shape corresponding to pre/post peak flow and point color corresponding to days since peak flow), and latent biomass (green polygon). Grey points in Panels C, D, and E are all GPP data within the year, and predicted estimates of GPP (C & E), and latent biomass (D) are shown as black lines with point shape corresponding to pre/post peak flow and point color corresponding to days since peak flow.

## 4 Discussion

A model of river ecosystem productivity that incorporates underlying autotrophic population dynamics provides ecological insights into river biomass resilience and disturbance thresholds. Model coverage by predictions from the latent biomass model (LB-TS) was high across rivers, indicating the strong predictive power of external light and flow forcing on temporal patterns in river ecosystem productivity (Roberts *et al*., 2007; Hall *et al*., 2015; Savoy *et al*., 2019). Deviations from these model predictions suggest a stronger influence of dynamics internal to the river ecosystem such as shifts in grazing (Pringle & Hamazaki, 1997, 1998), community composition (Vadeboncoeur & Power, 2017; Jiménez *et al*., 2023), or light availability because of turbidity in the water column (Savoy & Harvey, 2021b; Kirk *et al*., 2021) that our model does not capture. Streams and rivers are among the most frequently disturbed ecosystems, which selects for resilient primary producers (Stanley *et al*., 2010). Our modeling approach showed fast recovery of autotrophic biomass following disturbance with daily maximum percent increases ranging from 5-42% (2-14 d doubling times). The estimated magnitude of flood necessary to disturb biomass and thereby reduce ecosystem productivity was always lower than the more commonly used disturbance flow threshold of the flood magnitude necessary to mobilize river bed sediment (i.e., 2-y flood) indicating a greater sensitivity of productivity to disturbance than previously assumed. The LB-TS model presented here provides a tool by which to investigate controls on river resilience and productivity disturbance thresholds without overfitting relative to simpler models which often make better predictions than more complex ones (Perretti *et al*., 2013; Ward *et al*., 2014; Currie, 2019).

### 4.1 River productivity resilience to disturbances

Rivers and streams are disturbance-prone ecosystems (Stanley *et al*., 2010; Junk *et al*., 1989; Resh *et al*., 1998), but benthic autotrophic organisms in rivers are largely adapted to be resilient within the bounds of the natural flow regime (Poff *et al*., 1997). While the estimated carry capacity (*K*) overlapped in most of the six rivers included in this study, the broad distribution of site-specific *r_max_* estimates suggests that the conditions that facilitate a higher river *K* do not necessarily also determine how quickly a river can recover from disturbances. The largest river (Pecos River, TX) had both the highest *K* and lowest *r_max_* suggesting that the recovery of autotrophic biomass following disturbance might be slower in larger rivers. And yet, no clear pattern emerged between the six river *r_max_* estimates and the covariates we examined; however, our sample size of rivers was small. Further exploration of controls on estimates of ecosystem productivity to resilience across a wide range of environmental conditions among many rivers may clarify controls on disturbance recovery rates.

GPP tracked light availability with a hysteresis pattern as a result of lagged latent biomass recovery rates, matching our conceptual understanding of the reasons underlying disturbance recovery responses in streams (O’Donnell & Hotchkiss, 2022; Uehlinger, 2000; Qasem *et al*., 2019). However, the LB-TS model tended to underpredict the rate and peak of the GPP recovery trajectory across sites in response to disturbances, particularly in the spring. These patterns suggest that factors internal to the river ecosystem apart from external forcing of surface light availability and flood disturbances can strongly influence the GPP dynamics. Biological interactions such as grazing may vary seasonally in how they control autotrophic biomass, and therefore GPP (Power *et al*., 2008; Vadeboncoeur & Power, 2017). During late summer, apart from grazing, senescence and sloughing of algae may also decrease GPP in a way unexplained by external forcing (Graba *et al*., 2014). In their estimates of the recovery of insect species in rivers following disturbances, McMullen *et al*. (2017) used the simplifying assumption that *r_max_* is relatively constant while the carrying capacity, *K* varied through time. It is possible that *K* varies in rivers through time, particularly in rivers with expansion and contraction of wetted area (Stanley *et al*., 1997; Ruffing *et al*., 2022), shifts in the distribution of the autotrophic community from benthic to planktonic (Genzoli & Hall, 2016), or shifts in the relative importance of top-down (e.g., omnivory, herbivory) versus bottom-up (e.g., flow disturbance) controls on productivity (Pringle & Hamazaki, 1998, 1997).

### 4.2 Productivity disturbance thresholds

The magnitude of flood necessary to disturb biomass and thereby reduce whole-ecosystem GPP (*Q_c_*) was consistently lower than the estimated 2-y recurrence interval flood magnitude (*Q*_2_*_yr_*) necessary to mobilize the entire bed of a river and disturb the benthos (Fig. 3; Wolman & Miller (1960)). However, uncertainty is associated with estimating disturbance flow thresholds from GPP data. First, the ability of the model to identify the disturbance threshold parameters requires observing variation in flow, and in particular, observing disturbance flows. For example, the Pecos River only had one disturbance per year, resulting in more uncertainty in and tighter covariance between *c* and *s* (Fig. S18). Our comparison of training data set length in the South Branch Potomac River demonstrated that while fitting the model to more years of data may have improved most parameter estimates, the addition of years of training data that were increasingly different from the out-of-sample GPP data in their magnitude of GPP and frequency of storms made the identification of the critical flow disturbance threshold *c* more challenging for the LB-TS model thereby increasing process error. Second, floods often increase turbidity, complicating the separation of these covarying factors (Hall *et al*., 2015; Kirk *et al*., 2021; Savoy & Harvey, 2021b; Deemer *et al*., 2022). Regardless, whether a flood physically removed benthic biomass or suppressed productivity through persistent elevated turbidity following a storm, the LB-TS model still captured what are likely the combined effects of physical disturbance and turbidity-caused light attenuation by estimating the flood threshold necessary to induce those combined effects and the recovery rates specific to each river.

### 4.3 Conclusions

The modeling framework presented here can estimate a proxy for ecosystem productivity resilience (*r_max_*) and a site-specific disturbance flow threshold (*Q_c_*), demonstrating the ecological insights that can be gained from latent autotrophic biomass dynamics underlying patterns in GPP time series. The rate of autotrophic biomass recovery was distinct from the carrying capacity of a river and disturbance thresholds of river productivity were consistently lower than the magnitude necessary to mobilize the river bed. Although increasing model complexity often amplifies uncertainty in predicting new data (Getz *et al*., 2018; Ward *et al*., 2014; Tredennick *et al*., 2017; Perretti *et al*., 2013), our latent biomass model retained predictive capability relative to a simpler model and provided targeted ecological insights (Rastetter, 2017; Tredennick *et al*., 2021). We did not—and could not—attempt to capture all of the complexity that governs river productivity dynamics (e.g., grazing, senescence, and sloughing). Different modeling approaches may be better suited to improve the accuracy of a GPP forecast for a specific location, such as detailed models fit directly to dissolved oxygen observations that include a full spectrum of mechanisms controlling both heterotrophic and autotrophic biomass (e.g., Segatto *et al*. (2020)). Yet, complex models developed for specific locations are often challenging to transfer to new locations without substantial model tuning. The appropriate modeling framework to understand carbon fluxes in river ecosystems depends on the primary purpose of the model, ranging from gaining inference from specific parameters to high prediction accuracy (Currie, 2019; Rastetter, 2017; Tredennick *et al*., 2021). A diversity of modeling approaches that expand beyond only incorporating light and flow as the dominant controls on river metabolism (Bernhardt *et al*., 2022) will advance our collective understanding of the ecological processes underlying carbon fluxes in river ecosystems and how these fluxes are responding to environmental change (Battin *et al*., 2023).

## Acknowledgements

This research was supported by the StreamPulse project with funding from the National Science Foundation’s Division of Environmental Biology (1834679) and the modelscape project with funding from NSF Office of Integrative Activities (2019528). We thank Joel Scheingross for helpful comments regarding geomorphology and participants in the StreamPulse project for help-ful comments on early iterations of this work. Thank you to Nick Marzolf, Kathi Jo Jankowski, three anonymous reviewers, and the editor for comments that greatly improved this manuscript. Any use of trade, firm, or product names is for descriptive purposes only and does not imply endorsement by the U.S. Government.

## Data accessibility

Data for this project are in Appling *et al*. (2018b), Savoy & Harvey (2021a), and Blaszczak *et al*. (2021). Code to replicate the analyses is available at https://github.com/jrblaszczak/RiverBiomass.

## 6. Tables

**Table 1:**
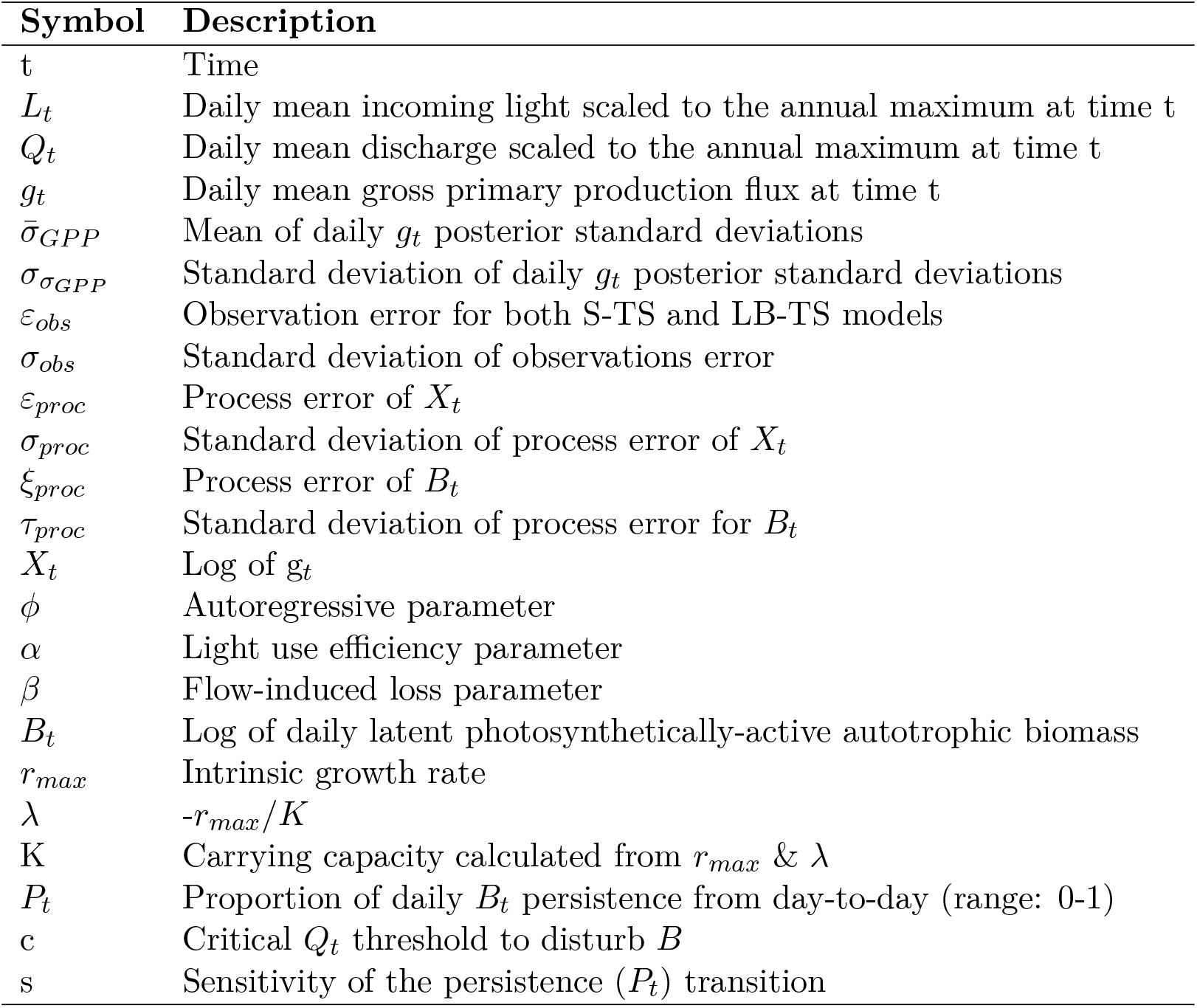
Variables and estimated parameters for the S-TS and LB-TS models.

| Symbol | Description |
| --- | --- |
| $t$ | Time |
| $L_t$ | Daily mean incoming light scaled to the annual maximum at time $t$ |
| $Q_t$ | Daily mean discharge scaled to the annual maximum at time $t$ |
| $g_t$ | Daily mean gross primary production flux at time $t$ |
| $\bar{\sigma}_{GPP}$ | Mean of daily $g_t$ posterior standard deviations |
| $\sigma_{\sigma_{GPP}}$ | Standard deviation of daily $g_t$ posterior standard deviations |
| $\varepsilon_{obs}$ | Observation error for both S-TS and LB-TS models |
| $\sigma_{obs}$ | Standard deviation of observations error |
| $\varepsilon_{proc}$ | Process error of $X_t$ |
| $\sigma_{proc}$ | Standard deviation of process error of $X_t$ |
| $\xi_{proc}$ | Process error of $B_t$ |
| $\tau_{proc}$ | Standard deviation of process error for $B_t$ |
| $X_t$ | Log of $g_t$ |
| $\phi$ | Autoregressive parameter |
| $\alpha$ | Light use efficiency parameter |
| $\beta$ | Flow-induced loss parameter |
| $B_t$ | Log of daily latent photosynthetically-active autotrophic biomass |
| $r_{max}$ | Intrinsic growth rate |
| $\lambda$ | $-r_{max}/K$ |
| $K$ | Carrying capacity calculated from $r_{max}$ & $\lambda$ |
| $P_t$ | Proportion of daily $B_t$ persistence from day-to-day (range: 0-1) |
| $c$ | Critical $Q_t$ threshold to disturb $B$ |
| $s$ | Sensitivity of the persistence ($P_t$) transition |

