## Appendix for "Models of underlying autotrophic biomass dynamics fit to daily river ecosystem productivity estimates improve understanding of ecosystem disturbance and resilience"

### A Supplemental Methods

#### A.1 Discrete time Ricker Model derivation

Start with the Ricker model:

$$N_t = N_{t-1}e^{r_{max}(1-\frac{N_{t-1}}{K})} \tag{S1}$$

Take the natural log of both sides of the equation:

$$ln(N_t) = ln(N_{t-1}) + r_{max}(1 - \frac{N_{t-1}}{K}) \tag{S2}$$

$$ln(N_t) = ln(N_{t-1}) + r_{max} - \frac{r_{max}}{K}N_{t-1} \tag{S3}$$

Substitute $B_t = ln(N_t)$ such that $B_t$ is the natural log of the latent biomass:

$$B_t = B_{t-1} + r_{max} - \frac{r_{max}}{K}e^{B_{t-1}} \tag{S4}$$

where $\lambda = -\frac{r_{max}}{K}$ in the final model.

#### A.2 Evolving information state calculation

We compared the relative predictive abilities of both the standard time series model (abbreviated as $S$ below) and the latent biomass time series model (abbreviated as $L$ below) by comparing their model weights through calculating the evolving information state of each model over the predicted out-of-sample year for each site to estimate which model accumulated the most supporting evidence (Nichols *et al.*, 2019, 2021). The evolving information state approach calculates a cumulative model weight, or relative confidence, for each time point based on the previous model weight and how well the mean and standard error of the model predictions predict the data. For each out-of-sample predicted time series of GPP, we first extracted the mean and standard error for each predicted GPP day for both the $S$ and $L$ models. We then calculated the probability of each each observation ($obs_t$) occurring on day $t$ given each model prediction (i.e., $P(obs_t|S)$ and $P(obs_t|L)$) using the 'dnorm' function in R. We began each evolving information state comparison with an even split of confidence (0.5 each) between models, and subsequent model weights updated with each new set of observations according to:

$$W_{S,t} = \frac{W_{S,t} \times P(obs_t|S)}{(W_{S,t-1} \times P(obs_t|S)) + (W_{L,t-1} \times P(obs_t|L)))} \tag{S5}$$

$$W_{L,t} = 1 - W_{S,t} \tag{S6}$$

where $W_{S,t}$ is the weight towards the $S$ model after each new observation, and $W_{S,t-1}$ is the model weight before the observation, $P(obs_t|S)$ is likelihood (deviance) of each new observation under the $S$ model, and similar definitions for the $L$ model components. At each step, model weights sum to 1 and as evidence accumulates in support of one model over the other, the associated weight for the more successful model approaches 1 and the associated weight for the less successful model approaches 0. Particularly strong deviations in a competing model's predictions from the data will cause a model to lose weight rapidly relative to the other model(s) it is being compared against, which can lead to alternating model weights throughout a year of GPP predictions (Fig. S11).

##### A.3 Flood exceedance probability calculation

To calculate the exceedance probabilities for floods of a given discharge we followed the procedure described in Hornberger *et al.* (1998). We downloaded USGS mean daily discharge data for each site from January 1, 1970 to December 31, 2020. For each site, we first determined the maximum daily discharge per year for each site and then ranked years by the magnitude of the maximum daily discharge with the largest flood assigned a rank of 1 and declining integers for subsequently smaller maximum flood magnitudes per year. We calculated the exceedance probability by calculating the fraction of floods greater than or equal to that flood by dividing the rank of a particular flood ($r$) by the number of total years in the record ($n$) plus one (i.e. $r/(n+1)$). We then fit a linear relationship between the log of the maximum annual discharge and the corresponding exceedance probability using maximum likelihood estimation, and used the relationship to estimate the magnitude of the 2 y flood recurrence interval discharge ($Q_{2yr}$) as,

$$Q_{2yr} = e^{\frac{((\frac{1}{2})-\beta)}{\alpha}} \tag{S7}$$

where $\alpha$ is the slope and $\beta$ is the intercept of the linear relationship between the log of the maximum annual discharge and the corresponding exceedance probability.

### B Supplemental Figures

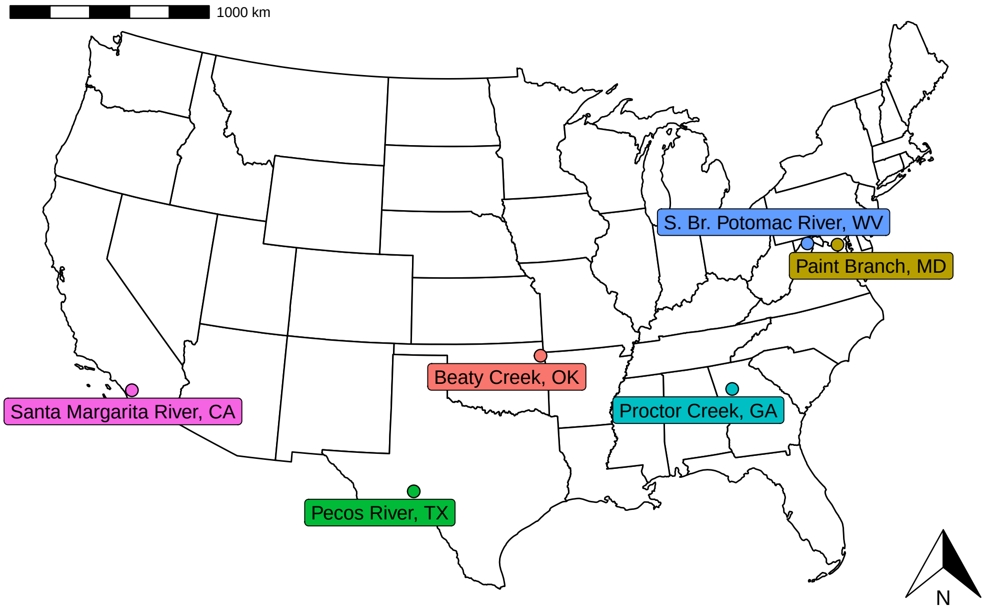


Figure S1: Map of six river locations included in this study.

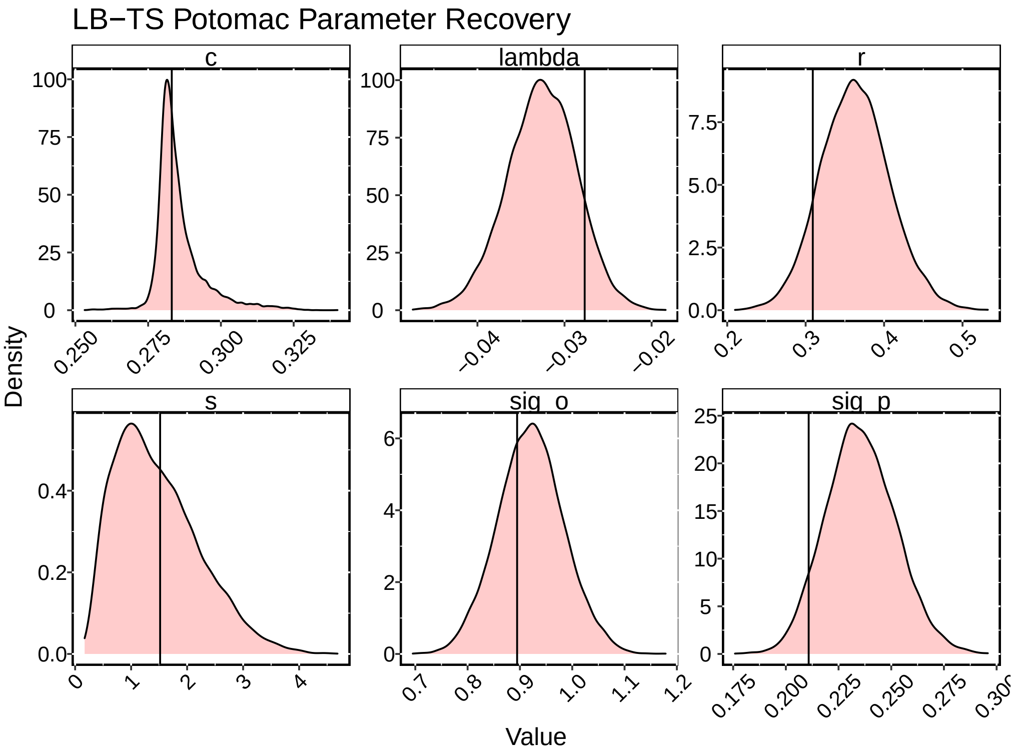


Figure S2: Parameter recovery for the LB-TS model fit to one year of simulated GPP data using the parameter values shown as black vertical lines and discharge and light data from the first year of data for the South Branch of the Potomac River, WV. Posterior distributions for each parameter are shown in pink.

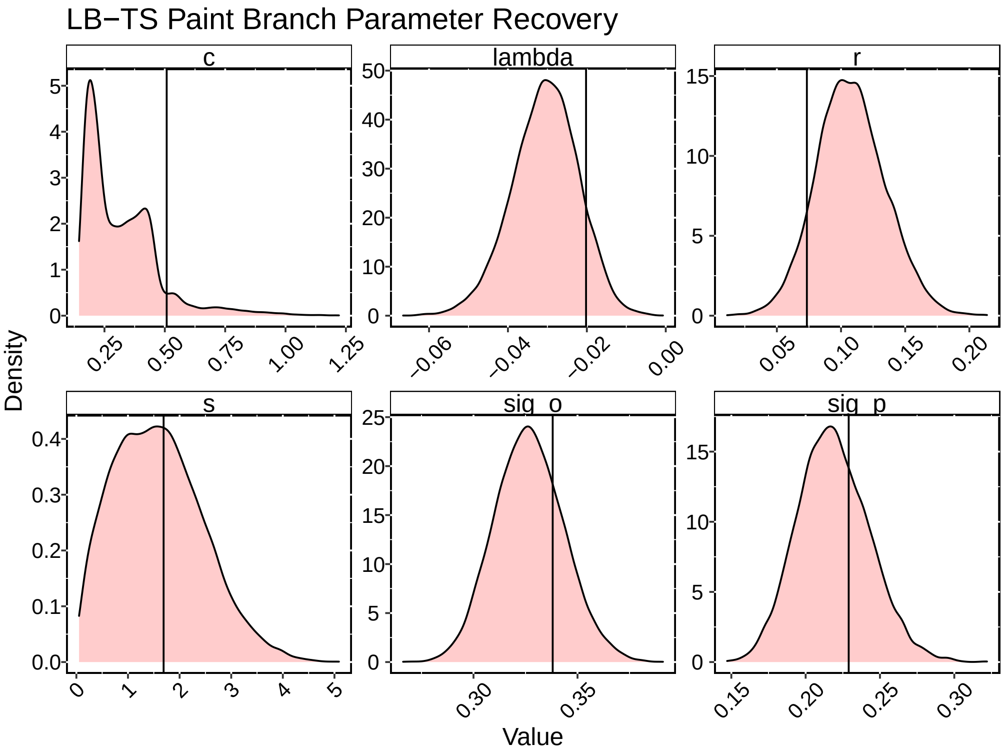


Figure S3: Parameter recovery for the LB-TS model fit to one year of simulated GPP data using the parameter values shown as black vertical lines and discharge and light data from the first year of data for the Paint Branch, MD. Posterior distributions for each parameter are shown in pink.

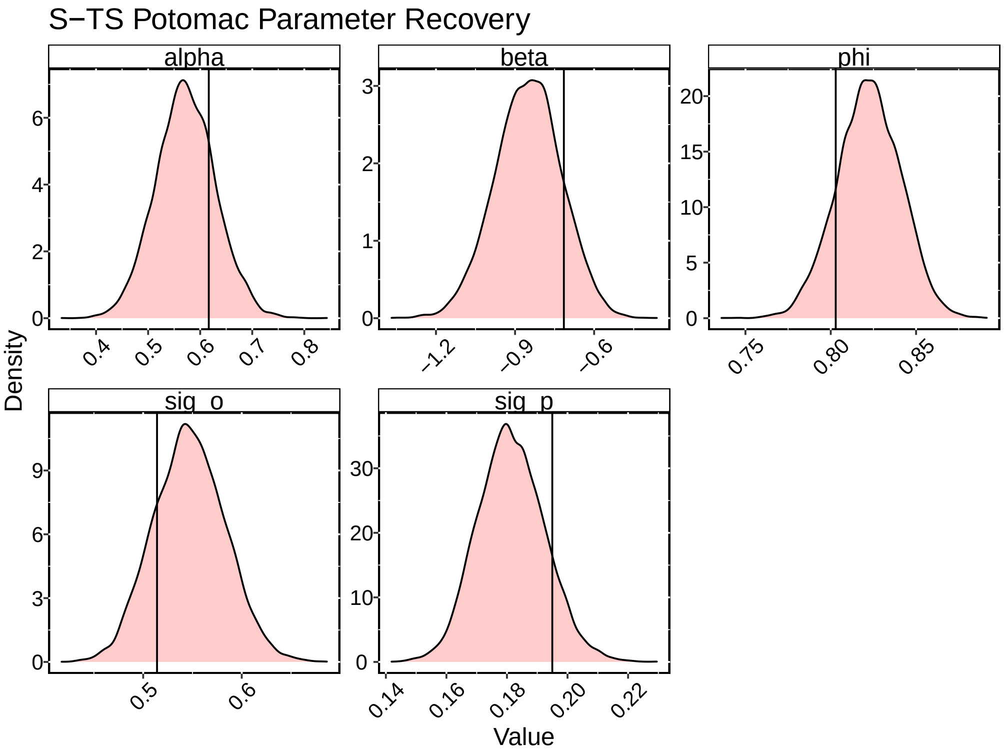


Figure S4: Parameter recovery for the S-TS model fit to one year of simulated GPP data using the parameter values shown as black vertical lines and discharge and light data from the first year of data for the South Branch of the Potomac River, WV. Posterior distributions for each parameter are shown in pink.

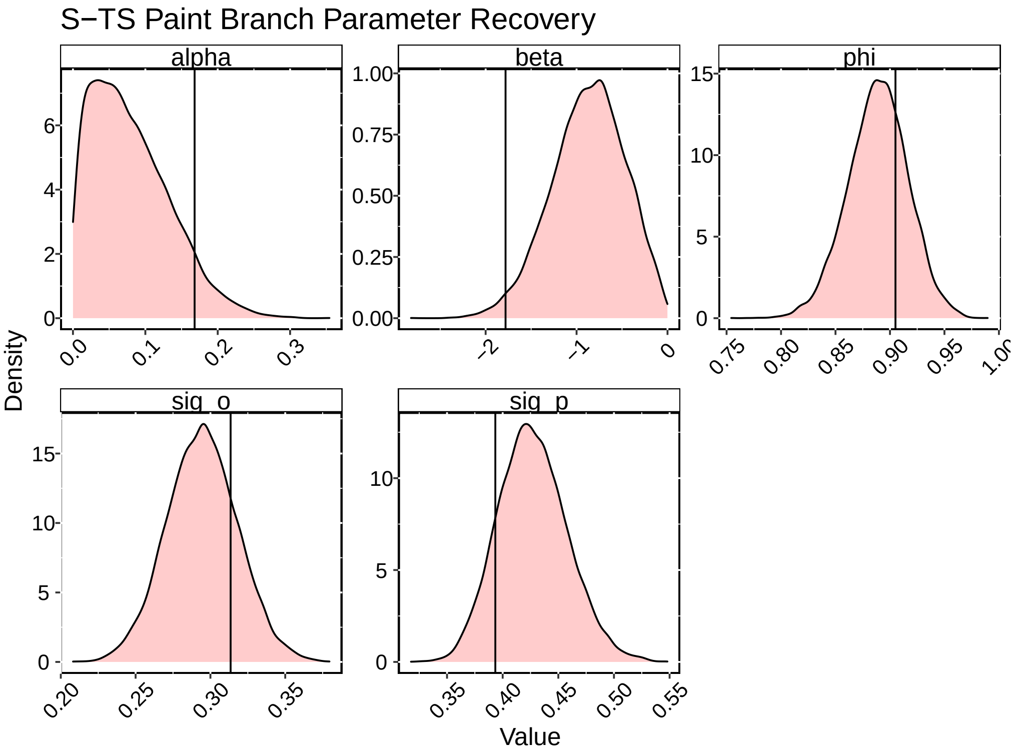


Figure S5: Parameter recovery for the S-TS model fit to one year of simulated GPP data using the parameter values shown as black vertical lines and discharge and light data from the first year of data for the Paint Branch, MD. Posterior distributions for each parameter are shown in pink.

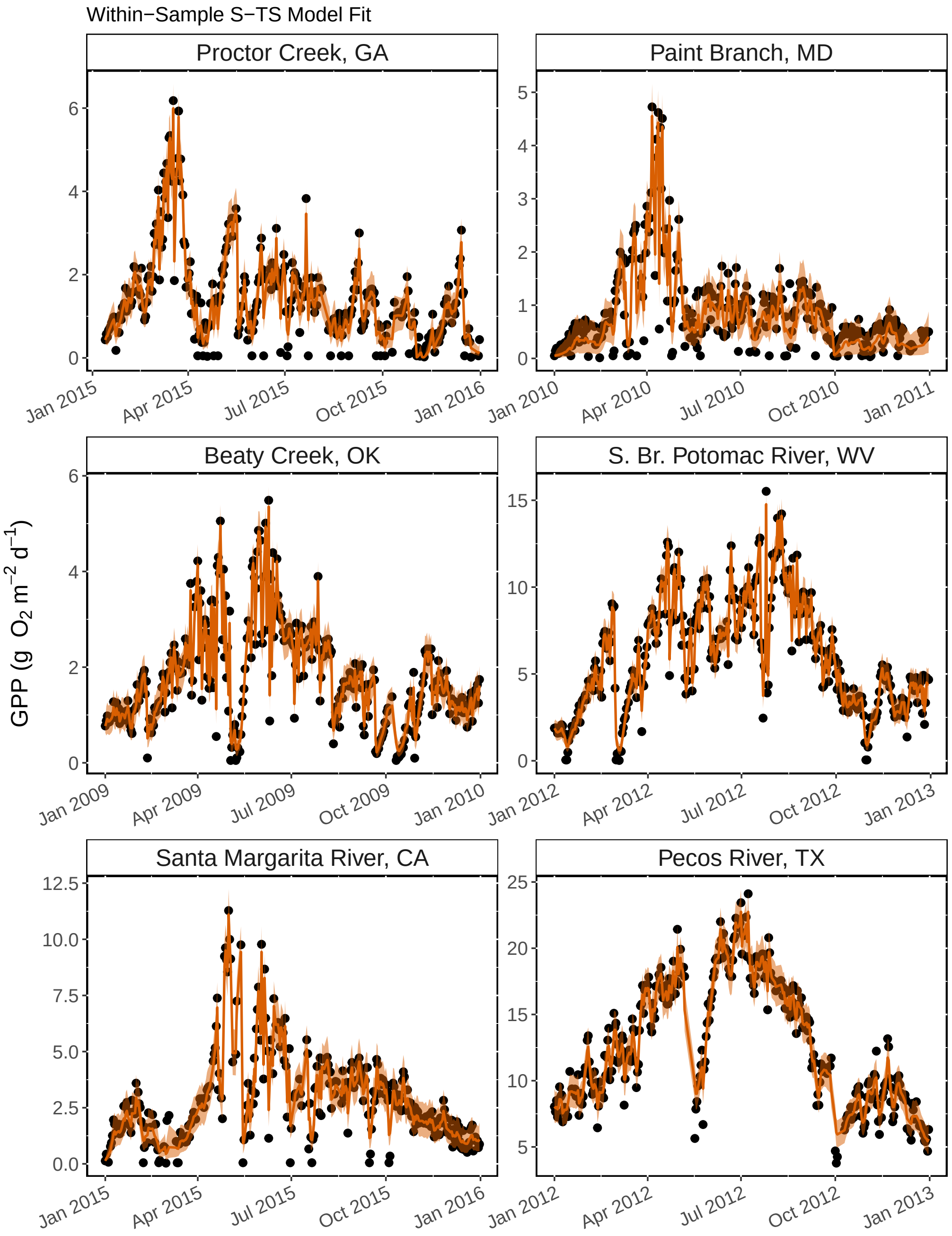


Figure S6: Median within-sample S-TS model estimates of daily GPP shown as orange lines with uncertainty intervals (2.5% - 97.5%) shown as light orange ribbons and original GPP data shown as black points for one year of data in each river included in this study.

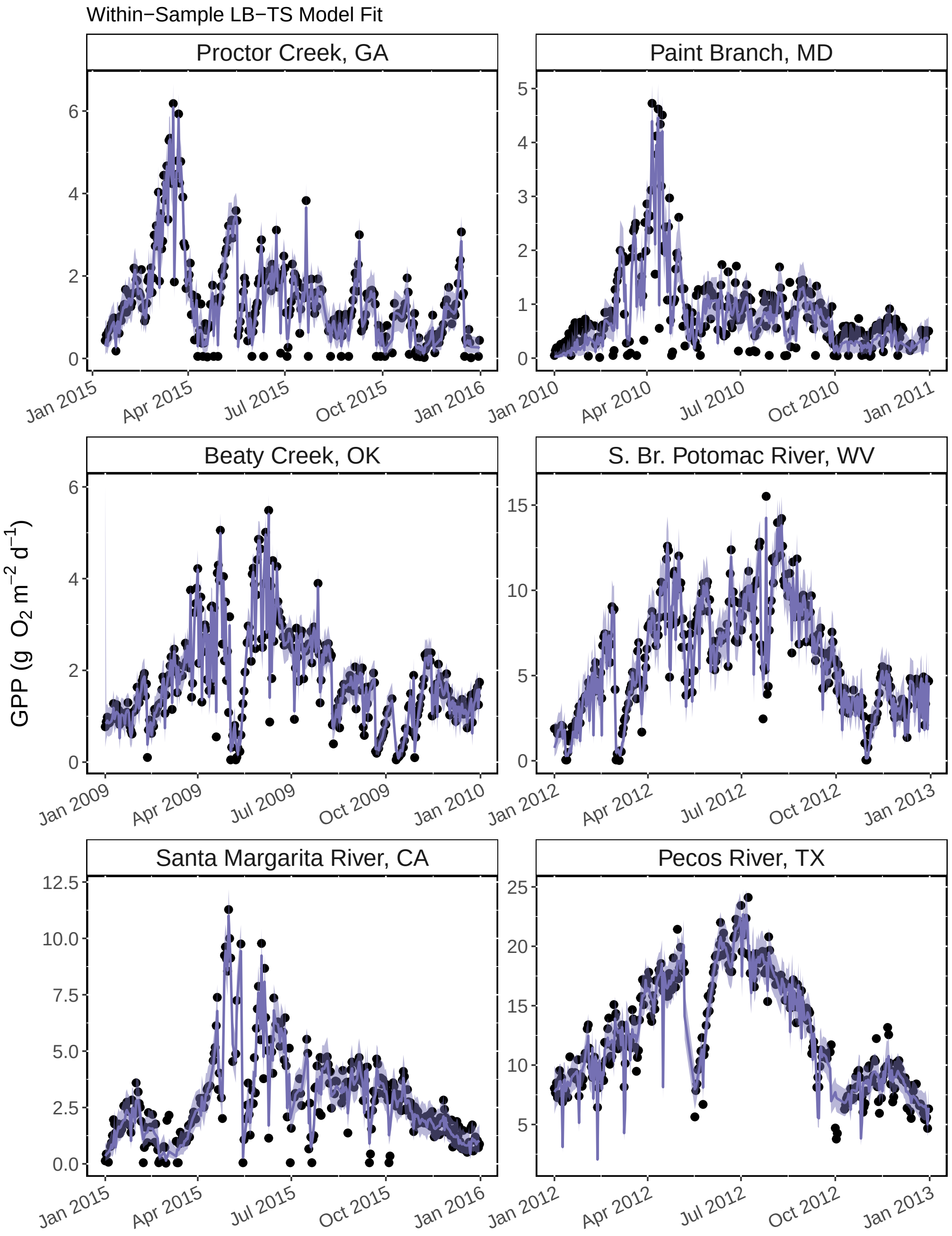


Figure S7: Median within-sample LB-TS model estimates of daily GPP shown as purple lines with uncertainty intervals (2.5% - 97.5%) shown as light purple ribbons and original GPP data shown as black points for one year of data in each river included in this study.

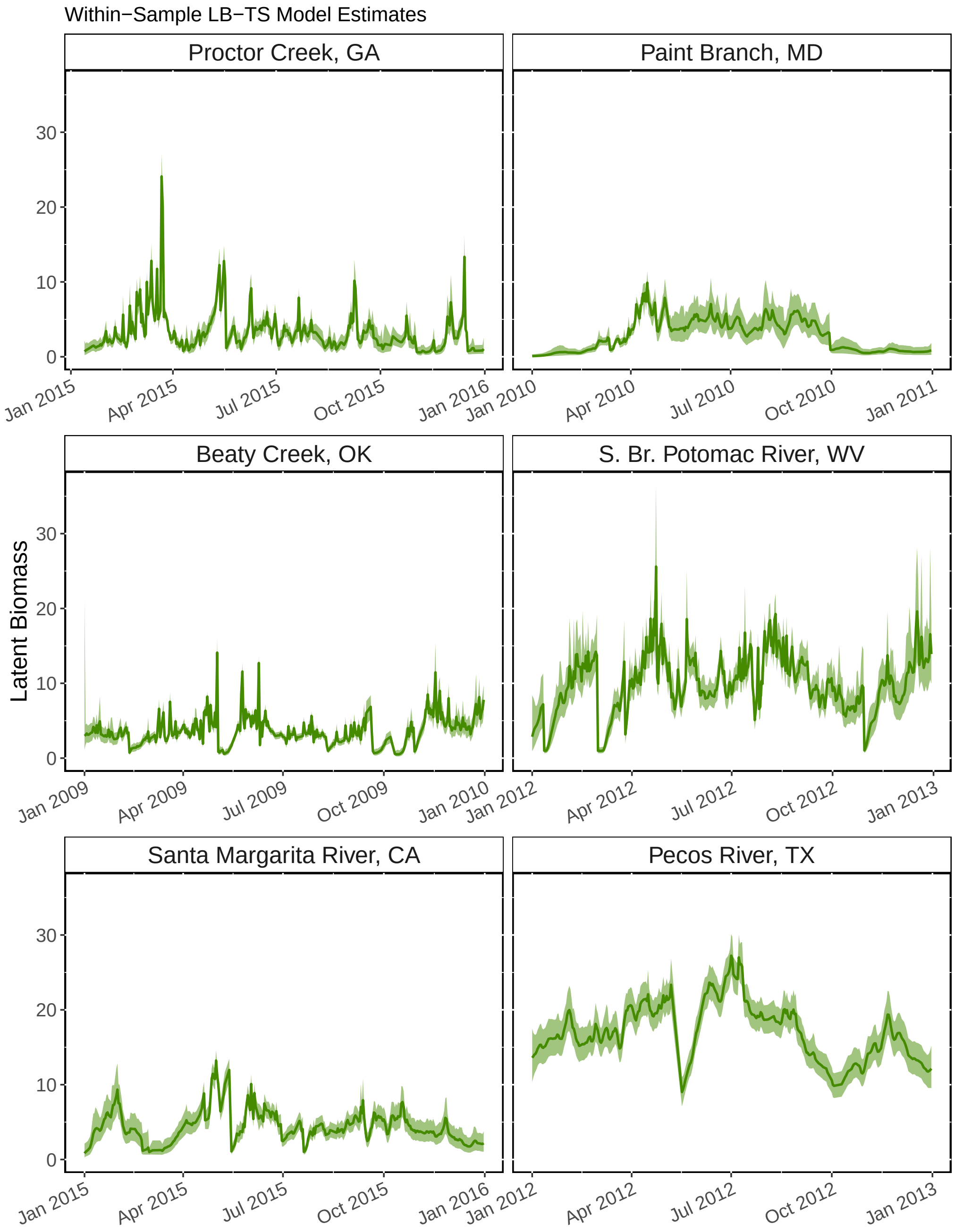


Figure S8: Median within-sample LB-TS model estimates of daily latent biomass shown as green lines with uncertainty intervals (2.5% - 97.5%) shown as light green ribbons for one year of data in each river included in this study and corresponding to the same output as in Figure S7.

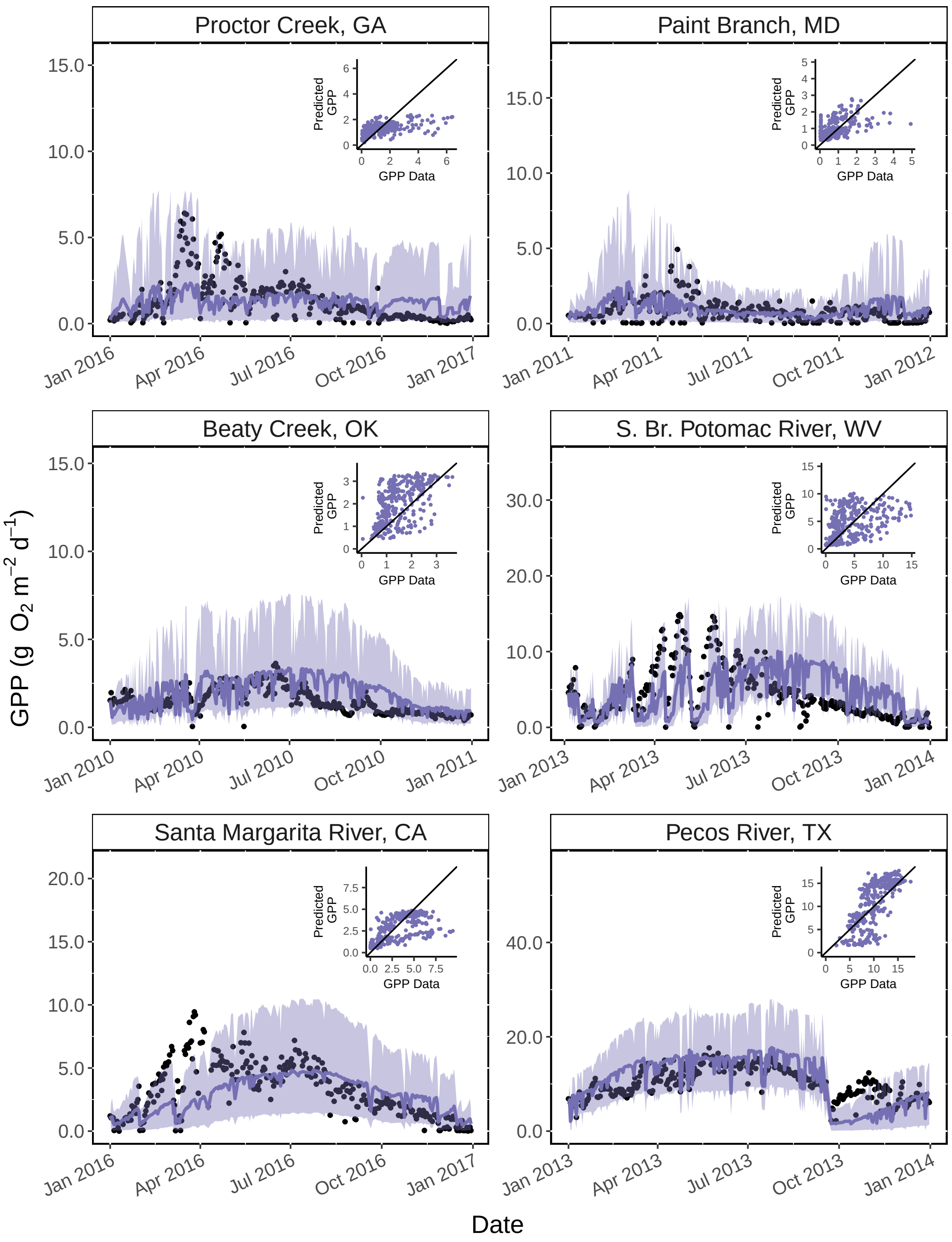


Figure S9: Out-of-sample predictions of daily gross primary productivity (GPP) over one year in six rivers using the latent biomass time series (LB-TS in purple) model. Black dots are previously estimated GPP data. Model predictions were generated using the full posterior distributions, and the median of the predictions are shown as lines and uncertainty intervals (2.5% - 97.5%) are shown as ribbons. Plots inset in each panel show the original GPP data on the x-axis and the LB-TS model predicted GPP on the y-axis.

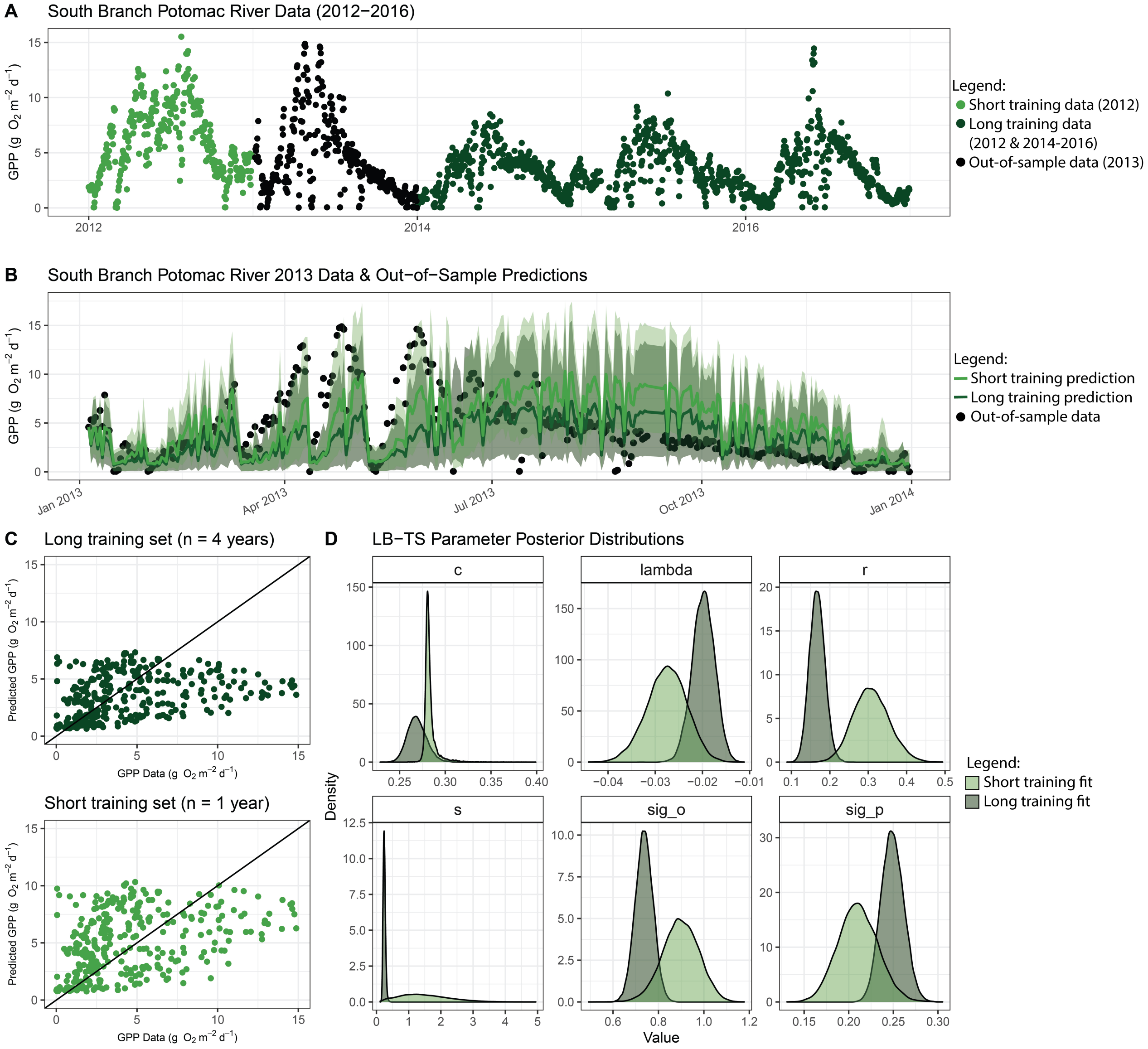


Figure S10: Comparison of how different training (i.e., within-sample) time series lengths influence LB-TS model out-of-sample predictions and parameter estimates. A) Original GPP time series for the South Branch Potomac River from 2012-2016 that met the criteria for inclusion in this study. Within-sample (2012) data originally used to fit (”train”) the LB-TS model is shown in light green points, out-of-sample (2013) data in shown in black points, and the longer training data which includes 2012 and also 2014, 2015, and 2016 is shown in light and dark green points. 2012 and 2013 are the most similar to each other in terms of the magnitude and variability in GPP. B) Median out-of-sample (2013) predictions using the full within-sample posterior parameter distributions for the short and long time series LB-TS model with uncertainty intervals (2.5% - 97.5%). Predictions based on short (2012) data model fit capture GPP peaks in the spring better while overestimating the late summer/fall GPP more than the longer data model fit. C) 1:1 comparisons of the predicted vs ”observed” GPP data for the long versus short training data. D) Density plots of the posterior parameter distributions for each model. A longer training data set results in more constrained parameter estimates for all parameters except $c$ which has a wider distribution likely because the GPP storm responses aren’t as clear in 2014-2016 as they are in 2012.

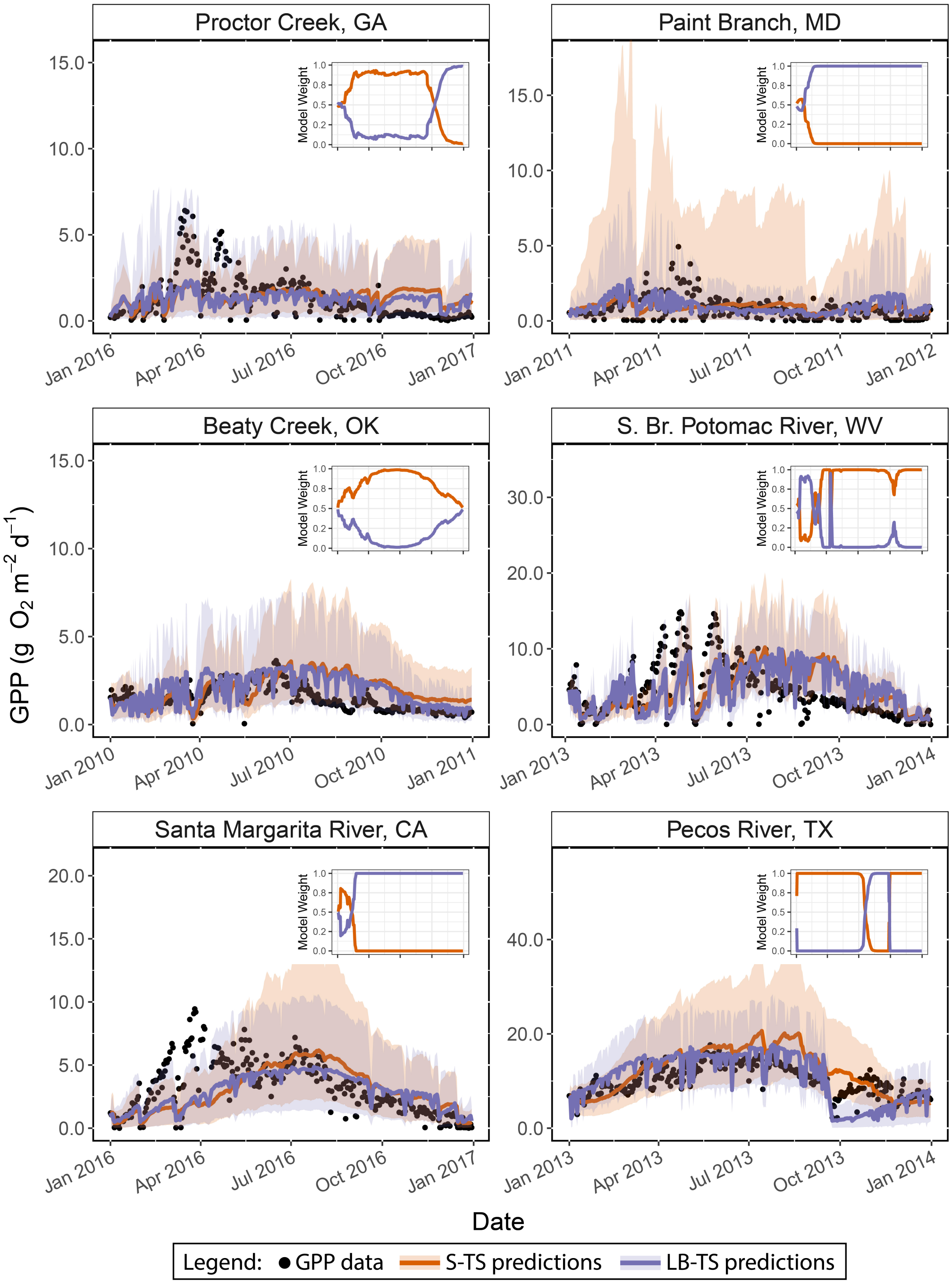


Figure S11: Out-of-sample predictions of daily gross primary productivity (GPP) over one year in six rivers using the standard time series (S-TS in orange) and latent biomass time series (LB-TS in purple) models. Black dots are previously estimated GPP data. Model predictions were generated using the full posterior distributions, and the median of the predictions are shown as lines and uncertainty intervals (2.5% - 97.5%) are shown as ribbons. Plots inset in each panel show which model accumulated the most support over the year of predictions, with the most successful model reaching a value of 1 by the end of the year.

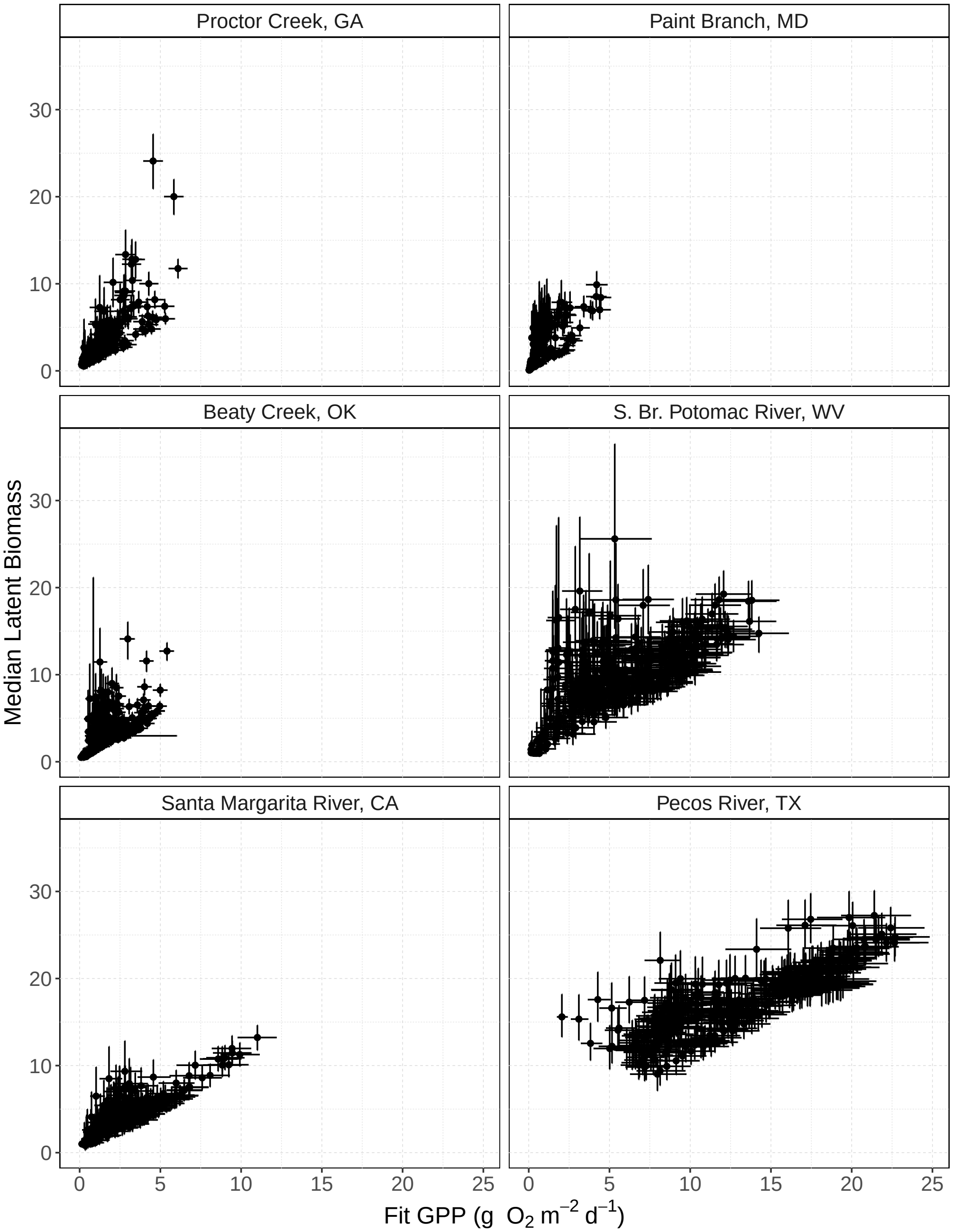


Figure S12: Within-sample covariance between median LB-TS model estimates of daily gross primary production (GPP; g $O_2$ $m^{-2}$ $d^{-1}$) and median latent biomass estimates with uncertainty intervals (2.5% - 97.5%) for each river location.

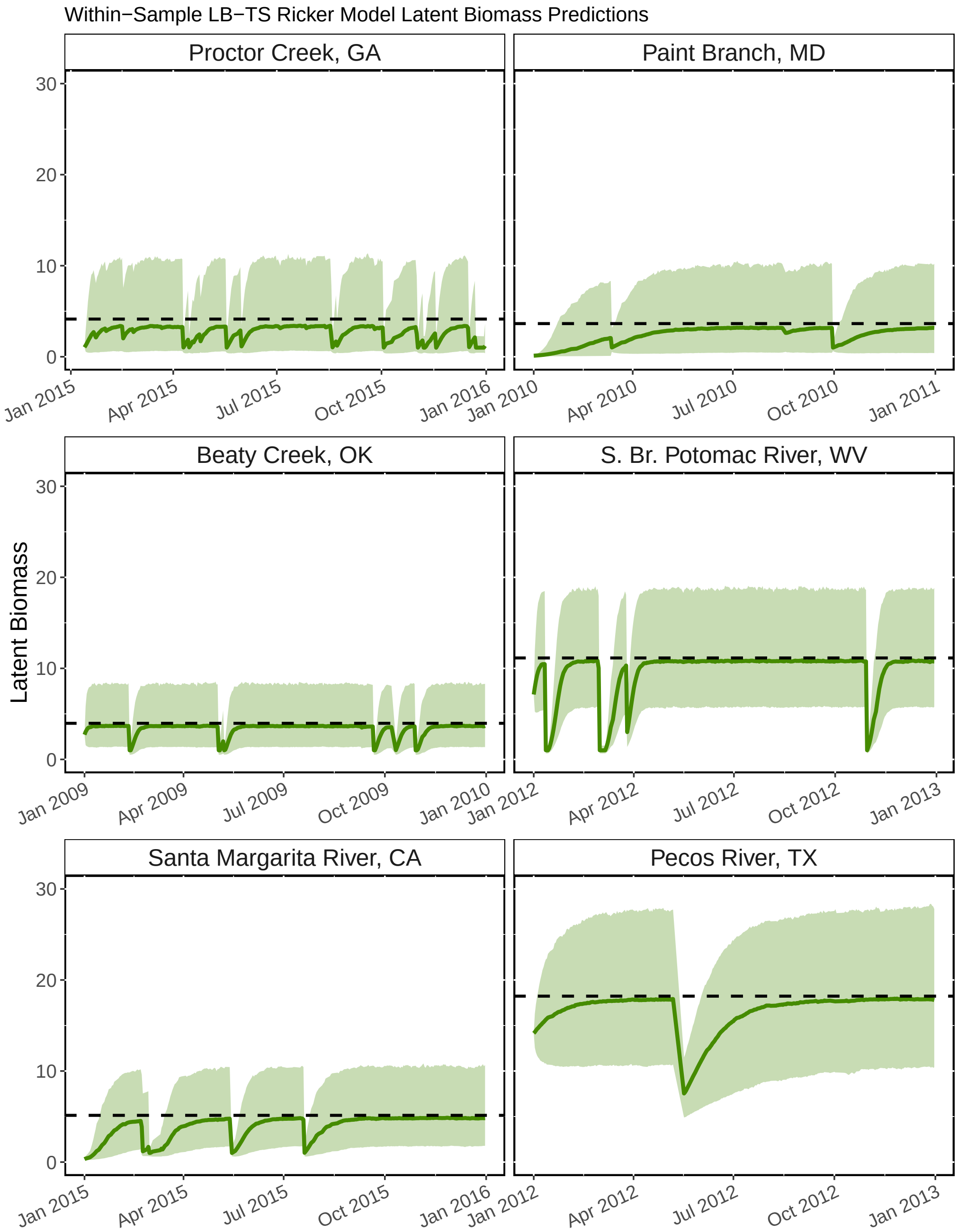


Figure S13: Latent biomass predictions generated using the full posterior parameter distributions from the within-sample LB-TS model fit shown in Figures S7 & S8. Median predictions are shown as green lines with uncertainty intervals (2.5% - 97.5%) shown as light green ribbons. Horizontal black dashed lines show the median estimated carrying capacity (K) for each river.

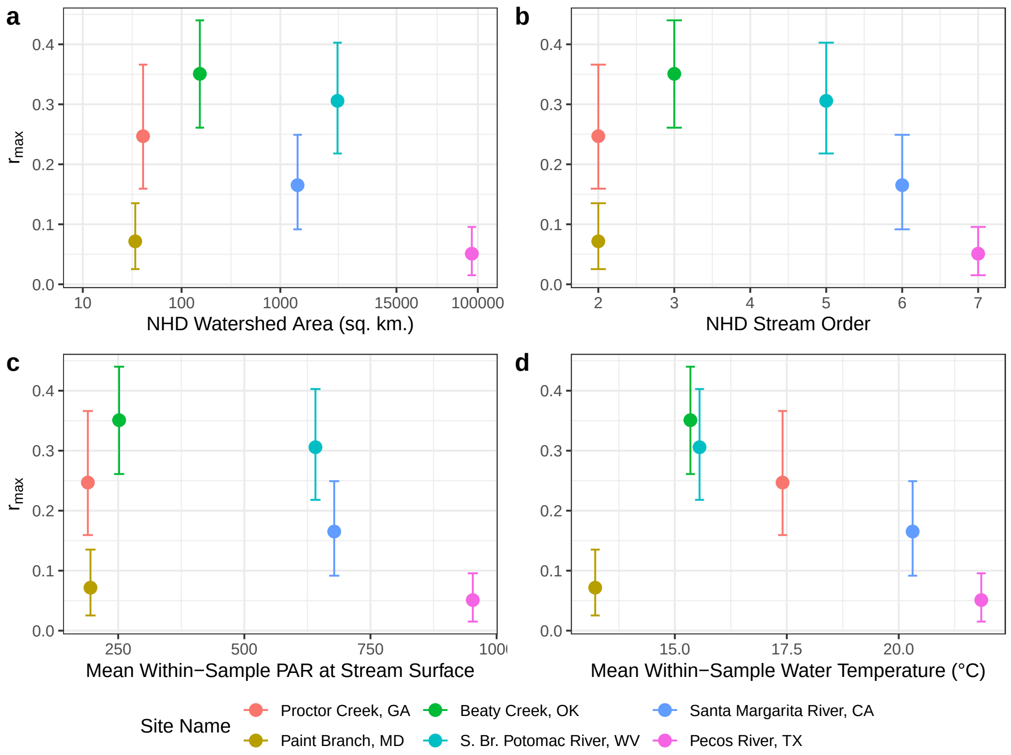


Figure S14: Within-sample median $r_{max}$ estimates with uncertainty intervals (2.5% - 97.5%) across a) NHDv2 watershed area (sq. km.), b) NHDv2 Strahler stream order designation, c) mean within-sample photosynthetically active radiation (PAR; $\mu$mol m$^{-2}$ s$^{-1}$) at the river surface, and d) the mean within-sample water temperature (°C).

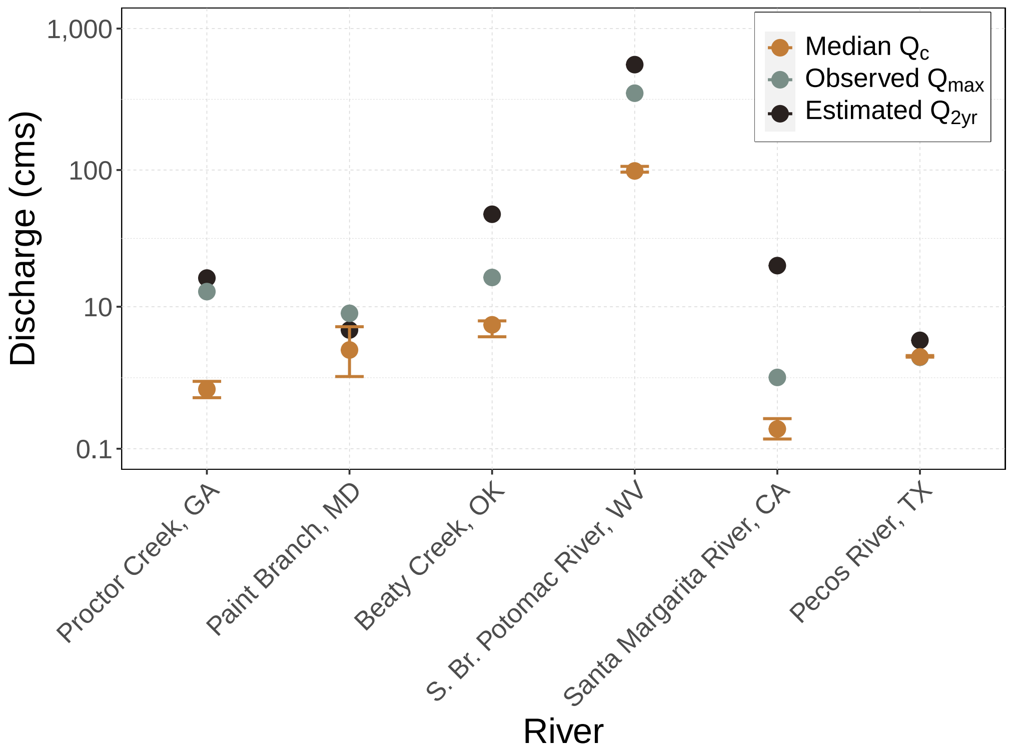


Figure S15: Comparison of within-sample discharge (cubic meters per second (cms)) disturbance thresholds for each river site including the observed maximum daily discharge (observed $Q_{max}$ in green-grey), the estimated two year recurrence interval flood (estimated $Q_{2yr}$ in black), and the median critical latent biomass disturbance threshold estimate with uncertainty intervals (2.5% - 97.5%; $Q_c$ in orange).

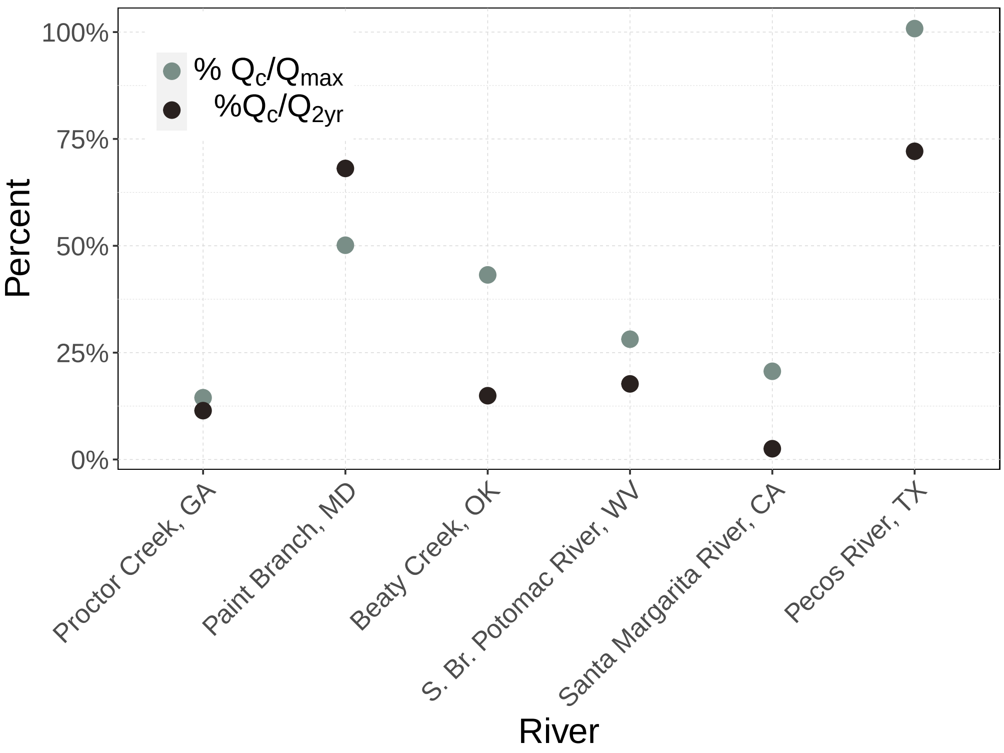


Figure S16: The relative magnitude of the median critical latent biomass disturbance threshold estimate ($Q_c$) as compared to the observed maximum daily discharge ($Q_{max}$) in green-grey, and compared to the estimated two year recurrence interval flood ($Q_{2yr}$) in black. The relative magnitude of nearly all $Q_c$ values are less than $Q_{max}$ or $Q_{2yr}$ apart from the relative magnitude of $Q_c$ to $Q_{max}$ in the Pecos River, TX.

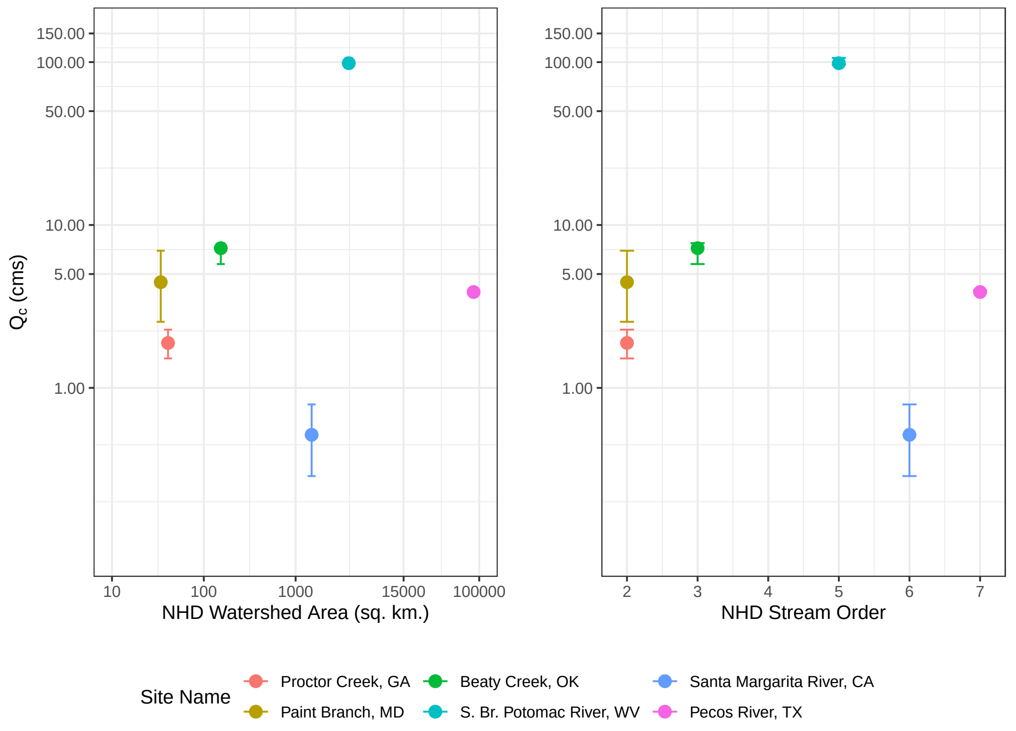


Figure S17: The median critical latent biomass disturbance threshold estimate ($Q_c$) with uncertainty intervals (2.5% - 97.5%) for each river location as compared to the contributing watershed area (sq. km.; left panel) and the NHDv2 Strahler stream order designation.

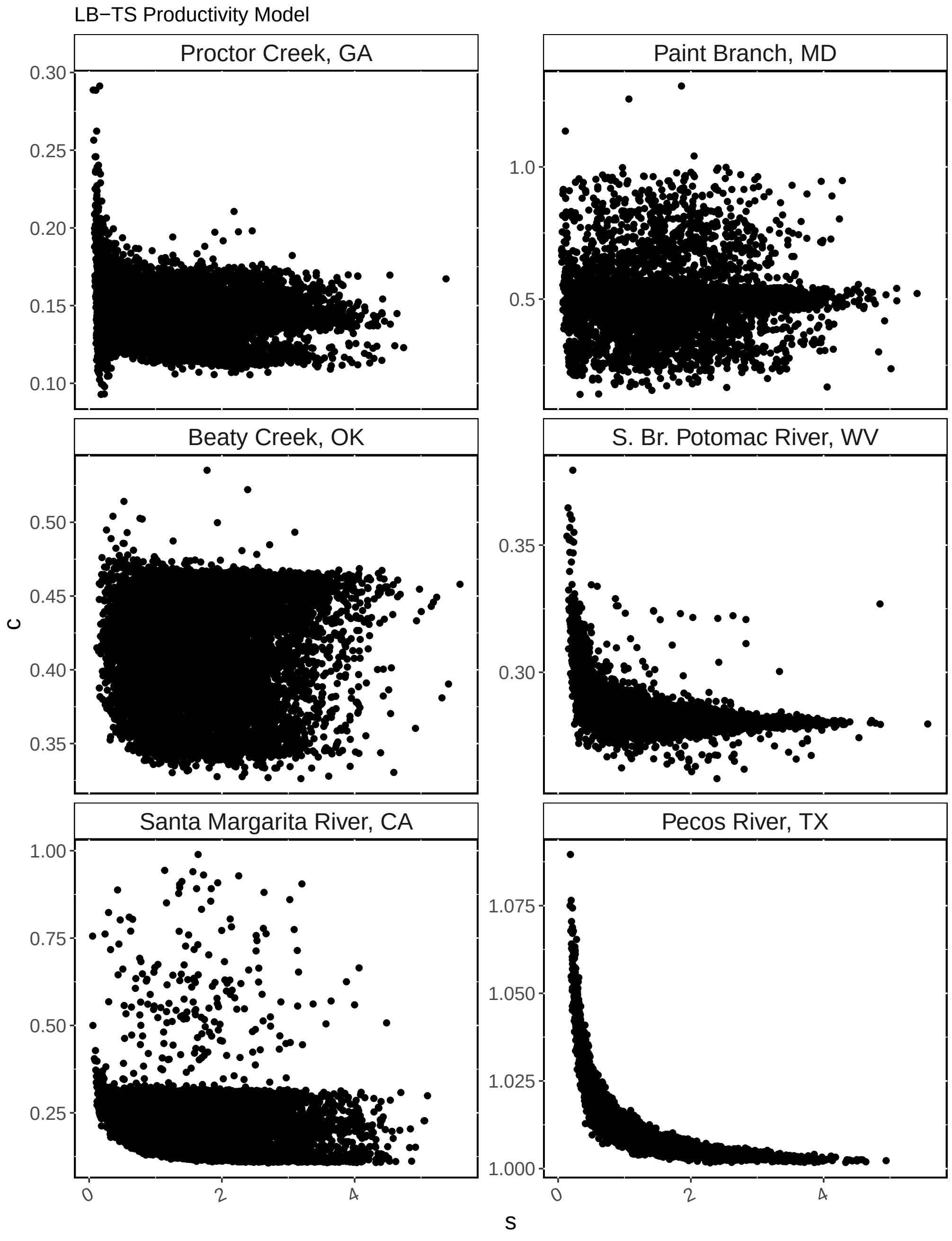


Figure S18: Covariance between every saved iteration of within-sample LB-TS model parameter estimates for the critical standardized flow threshold to disturb latent biomass, c, and the sensitivity of day-to-day latent biomass persistence from 0 to 1, s.

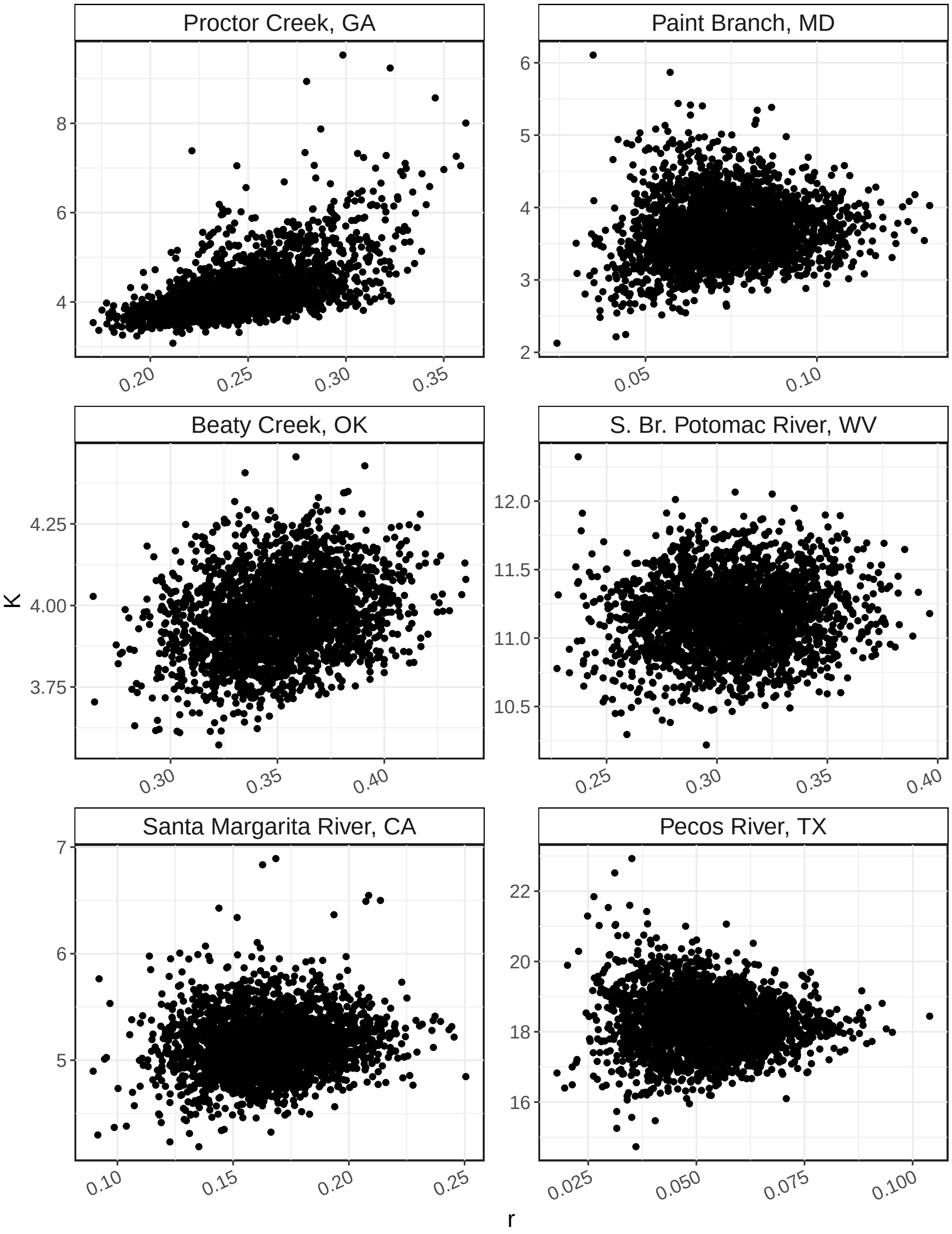


Figure S19: Covariance between every saved iteration of within-sample LB-TS model parameter estimates for the maximum growth rate ($r_{max}$) and the carrying capacity ($K$).

#### C Supplemental Tables

Table S1: Site information for each of the six river reach locations included in this study. Latitude and longitude coordinates correspond to the location of the U.S. Geological Survey maintained dissolved oxygen sensor that generated the raw data used to estimate metabolism. Stream Strahler order classification is from the StreamCat database (Hill *et al.*, 2016). Numeric flags for canal, dam, or National Pollution Discharge Elimination System (NPDES) point sources represent the percentiles of the mean daily 80% oxygen turnover distances that could be affected by the various upstream features which could affect metabolism estimates (Appling *et al.*, 2018b). Site flags indicate whether the nearest feature of each type was farther than the 95th, 80th, 50th, or 0th percentile. A value of 95 indicates the least probable interference from a structure of a given type.

| **Site Name** | **NWIS ID** | **Latitude** | **Longitude** | **Strahler Order** | **Canal Flag** | **Dam Flag** | **NPDES flag** |
|---|---|---|---|---|---|---|---|
| S. Br. Potomac River, WV | 1608500 | 39.44704 | -78.65418 | 5 | 95 | 95 | 80 |
| Paint Branch, MD | 1649190 | 39.03314 | -76.96428 | 2 | 95 | 95 | 50 |
| Proctor Creek, GA | 2336526 | 33.79427 | -84.47437 | 2 | 95 | 95 | 95 |
| Beaty Creek, OK | 7191222 | 36.35536 | -94.77634 | 3 | 95 | 95 | 95 |
| Pecos River, TX | 8447300 | 30.31400 | -101.74156 | 7 | 95 | 95 | 95 |
| Santa Margarita River, CA | 11044000 | 33.47392 | -117.14225 | 6 | 95 | 95 | 95 |

Table S2: Model diagnostic and coverage of daily modeled estimates of gross primary productivity (GPP) per year and site. Dissolved oxygen (DO) resolution in minutes corresponds to the original resolution of the DO data set that was used to generate the GPP estimates. The model observation and process error ($\sigma_{obs,iid}$ and $\sigma_{proc,iid}$, respectively) $\hat{R}$ values for the streamMetabolizer model that generated the GPP estimates are all less than 1.05. The maximum gap in continuous daily GPP estimates for each site within each year is less than 14 days, and the number of days of GPP per modeled year is greater than 275 days (75% of days in a year).

| **Site Name** | **DO resolution (min)** | **Full model** $\sigma_{obs,iid}$ $\hat{R}$ | **Full model** $\sigma_{proc,iid}$ $\hat{R}$ | **Year** | **Max gap (days) within year** | **Days in year** |
|---|---|---|---|---|---|---|
| S. Br. Potomac River, WV | 30 | 1.024 | 1.006 | 2012 | 2 | 352 |
| | | | | 2013 | 7 | 338 |
| Paint Branch, MD | 15 | 1.030 | 1.032 | 2010 | 8 | 312 |
| | | | | 2011 | 5 | 316 |
| Proctor Creek, GA | 15 | 1.008 | 1.001 | 2015 | 7 | 311 |
| | | | | 2016 | 7 | 323 |
| Beaty Creek, OK | 30 | 1.039 | 1.010 | 2009 | 4 | 341 |
| | | | | 2010 | 3 | 346 |
| Pecos River, TX | 15 | 1.005 | 1.007 | 2012 | 10 | 320 |
| | | | | 2013 | 5 | 337 |
| Santa Margarita River, CA | 15 | 1.010 | 1.010 | 2015 | 7 | 292 |
| | | | | 2016 | 5 | 282 |

Table S3: Prior specifications for both the standard time series (S-TS) and the latent biomass time series (LB-TS) models. All distributions are normal or truncated normal.

| **Model** | **Parameter** | **Mean** | **SD** | **Lower** | **Upper** | **Initial** |
|---|---|---|---|---|---|---|
| S-TS | $\phi$ | 0 | 1 | 0 | $\infty$ | |
| | $\alpha$ | 0 | 1 | 0 | $\infty$ | |
| | $\beta$ | 0 | 1 | $\infty$ | 0 | |
| | $\sigma_{obs}$ | mean $\sigma_{GPP}$ | SD $\sigma_{GPP}$ | 0 | $\infty$ | |
| | $\sigma_{proc}$ | 0 | 2 | 0 | $\infty$ | |
| LB-TS | $r_{max}$ | 0 | 1 | $\infty$ | $\infty$ | |
| | $\lambda$ | 0 | 1 | $\infty$ | 0 | |
| | s | 0 | 200 | 0 | $\infty$ | 100 |
| | c | 0 | 1 | 0 | $\infty$ | 0.5 |
| | $\sigma_{obs}$ | mean $\sigma_{GPP}$ | SD $\sigma_{GPP}$ | 0 | $\infty$ | |
| | $\sigma_{proc}$ | 0 | 2 | 0 | $\infty$ | |

Table S4: Within-sample parameter estimates from LB-TS model for the first year of each two year time series.

| | | Year 1 | | | | | |
|---|---|---|---|---|---|---|---|
| **Site Name** | **Parameter** | **50%** | **2.5%** | **97.5%** | $\hat{R}$ | $N_{eff}$ | $N_{eff}$ **<10%** |
| S. Br. Potomac River, WV | $c$ | 0.281 | 0.276 | 0.303 | 1.003 | 702 | TRUE |
| S. Br. Potomac River, WV | $s$ | 1.435 | 0.334 | 3.252 | 1.001 | 3974 | FALSE |
| S. Br. Potomac River, WV | $r_{max}$ | 0.306 | 0.218 | 0.403 | 1.003 | 1458 | FALSE |
| S. Br. Potomac River, WV | $\lambda$ | -0.027 | -0.036 | -0.019 | 1.003 | 1521 | FALSE |
| S. Br. Potomac River, WV | $\sigma_{proc}$ | 0.209 | 0.166 | 0.255 | 1.004 | 850 | TRUE |
| S. Br. Potomac River, WV | $\sigma_{obs}$ | 0.900 | 0.739 | 1.053 | 1.004 | 892 | TRUE |
| Paint Branch, MD | $c$ | 0.501 | 0.286 | 0.782 | 1.004 | 1080 | FALSE |
| Paint Branch, MD | $s$ | 1.623 | 0.245 | 3.465 | 1.000 | 9863 | FALSE |
| Paint Branch, MD | $r_{max}$ | 0.072 | 0.025 | 0.135 | 1.002 | 1344 | FALSE |
| Paint Branch, MD | $\lambda$ | -0.020 | -0.037 | -0.007 | 1.002 | 1141 | FALSE |
| Paint Branch, MD | $\sigma_{proc}$ | 0.228 | 0.158 | 0.308 | 1.006 | 412 | TRUE |
| Paint Branch, MD | $\sigma_{obs}$ | 0.338 | 0.288 | 0.389 | 1.002 | 923 | TRUE |
| Proctor Creek, GA | $c$ | 0.145 | 0.116 | 0.175 | 1.000 | 2651 | FALSE |
| Proctor Creek, GA | $s$ | 1.276 | 0.149 | 3.366 | 1.001 | 3248 | FALSE |
| Proctor Creek, GA | $r_{max}$ | 0.247 | 0.159 | 0.366 | 1.001 | 2401 | FALSE |
| Proctor Creek, GA | $\lambda$ | -0.060 | -0.081 | -0.035 | 1.000 | 4121 | FALSE |
| Proctor Creek, GA | $\sigma_{proc}$ | 0.418 | 0.367 | 0.475 | 1.001 | 2461 | FALSE |
| Proctor Creek, GA | $\sigma_{obs}$ | 0.292 | 0.243 | 0.343 | 1.001 | 1258 | FALSE |
| Beaty Creek, OK | $c$ | 0.432 | 0.345 | 0.464 | 1.001 | 3008 | FALSE |
| Beaty Creek, OK | $s$ | 1.722 | 0.419 | 3.571 | 1.000 | 13747 | FALSE |
| Beaty Creek, OK | $r_{max}$ | 0.351 | 0.261 | 0.440 | 1.001 | 4039 | FALSE |
| Beaty Creek, OK | $\lambda$ | -0.088 | -0.111 | -0.066 | 1.001 | 4225 | FALSE |
| Beaty Creek, OK | $\sigma_{proc}$ | 0.333 | 0.294 | 0.373 | 1.001 | 1730 | FALSE |
| Beaty Creek, OK | $\sigma_{obs}$ | 0.214 | 0.157 | 0.279 | 1.005 | 563 | TRUE |
| Pecos River, TX | $c$ | 1.008 | 1.003 | 1.035 | 1.001 | 3342 | FALSE |
| Pecos River, TX | $s$ | 1.255 | 0.340 | 3.091 | 1.000 | 7027 | FALSE |
| Pecos River, TX | $r_{max}$ | 0.051 | 0.015 | 0.096 | 1.001 | 1729 | FALSE |
| Pecos River, TX | $\lambda$ | -0.003 | -0.005 | -0.001 | 1.001 | 1660 | FALSE |
| Pecos River, TX | $\sigma_{proc}$ | 0.064 | 0.050 | 0.084 | 1.006 | 549 | TRUE |
| Pecos River, TX | $\sigma_{obs}$ | 1.598 | 1.450 | 1.756 | 1.001 | 2557 | FALSE |
| Santa Margarita River, CA | $c$ | 0.206 | 0.115 | 0.317 | 1.021 | 215 | TRUE |
| Santa Margarita River, CA | $s$ | 1.736 | 0.391 | 3.589 | 1.001 | 6232 | FALSE |
| Santa Margarita River, CA | $r_{max}$ | 0.165 | 0.092 | 0.249 | 1.001 | 2079 | FALSE |
| Santa Margarita River, CA | $\lambda$ | -0.032 | -0.049 | -0.017 | 1.001 | 2218 | FALSE |
| Santa Margarita River, CA | $\sigma_{proc}$ | 0.228 | 0.187 | 0.275 | 1.003 | 1232 | FALSE |
| Santa Margarita River, CA | $\sigma_{obs}$ | 0.631 | 0.534 | 0.742 | 1.002 | 1724 | FALSE |

Table S5: Within-sample parameter estimates from S-TS model for the first year of each two year time series.

| | | **Year 1** | | | | | |
|---|---|---|---|---|---|---|---|
| **Site Name** | **Parameter** | **50%** | **2.5%** | **97.5%** | $\hat{R}$ | $N_{eff}$ | $N_{eff}$ **<10%** |
| S. Br. Potomac River, WV | $\phi$ | 0.803 | 0.760 | 0.844 | 1.000 | 4604 | FALSE |
| S. Br. Potomac River, WV | $\alpha$ | 0.617 | 0.493 | 0.745 | 1.000 | 4057 | FALSE |
| S. Br. Potomac River, WV | $\beta$ | -0.709 | -0.962 | -0.475 | 1.000 | 3737 | FALSE |
| S. Br. Potomac River, WV | $\sigma_{proc}$ | 0.195 | 0.174 | 0.218 | 1.001 | 2392 | FALSE |
| S. Br. Potomac River, WV | $\sigma_{obs}$ | 0.514 | 0.415 | 0.617 | 1.003 | 948 | TRUE |
| Paint Branch, MD | $\phi$ | 0.908 | 0.835 | 0.966 | 1.002 | 1127 | FALSE |
| Paint Branch, MD | $\alpha$ | 0.165 | 0.029 | 0.324 | 1.000 | 4410 | FALSE |
| Paint Branch, MD | $\beta$ | -1.772 | -2.828 | -0.830 | 1.001 | 3090 | FALSE |
| Paint Branch, MD | $\sigma_{proc}$ | 0.392 | 0.318 | 0.474 | 1.001 | 1244 | FALSE |
| Paint Branch, MD | $\sigma_{obs}$ | 0.313 | 0.258 | 0.379 | 1.003 | 818 | TRUE |
| Proctor Creek, GA | $\phi$ | 0.798 | 0.742 | 0.851 | 1.001 | 2447 | FALSE |
| Proctor Creek, GA | $\alpha$ | 0.334 | 0.239 | 0.434 | 1.000 | 3379 | FALSE |
| Proctor Creek, GA | $\beta$ | -3.881 | -4.867 | -2.919 | 1.001 | 2068 | FALSE |
| Proctor Creek, GA | $\sigma_{proc}$ | 0.286 | 0.245 | 0.332 | 1.002 | 2135 | FALSE |
| Proctor Creek, GA | $\sigma_{obs}$ | 0.297 | 0.258 | 0.339 | 1.001 | 2185 | FALSE |
| Beaty Creek, OK | $\phi$ | 0.692 | 0.614 | 0.768 | 1.001 | 2778 | FALSE |
| Beaty Creek, OK | $\alpha$ | 0.467 | 0.358 | 0.582 | 1.001 | 2400 | FALSE |
| Beaty Creek, OK | $\beta$ | -1.266 | -1.619 | -0.938 | 1.001 | 2244 | FALSE |
| Beaty Creek, OK | $\sigma_{proc}$ | 0.297 | 0.260 | 0.336 | 1.001 | 1550 | FALSE |
| Beaty Creek, OK | $\sigma_{obs}$ | 0.216 | 0.164 | 0.274 | 1.005 | 616 | TRUE |
| Pecos River, TX | $\phi$ | 0.941 | 0.908 | 0.974 | 1.000 | 4047 | FALSE |
| Pecos River, TX | $\alpha$ | 0.208 | 0.099 | 0.320 | 1.000 | 3960 | FALSE |
| Pecos River, TX | $\beta$ | -0.011 | -0.047 | 0.000 | 1.000 | 14104 | FALSE |
| Pecos River, TX | $\sigma_{proc}$ | 0.089 | 0.074 | 0.105 | 1.001 | 1690 | FALSE |
| Pecos River, TX | $\sigma_{obs}$ | 0.991 | 0.826 | 1.153 | 1.002 | 1834 | FALSE |
| Santa Margarita River, CA | $\phi$ | 0.729 | 0.635 | 0.811 | 1.001 | 2415 | FALSE |
| Santa Margarita River, CA | $\alpha$ | 0.586 | 0.428 | 0.758 | 1.000 | 2809 | FALSE |
| Santa Margarita River, CA | $\beta$ | -1.649 | -2.353 | -1.033 | 1.000 | 5492 | FALSE |
| Santa Margarita River, CA | $\sigma_{proc}$ | 0.286 | 0.243 | 0.336 | 1.000 | 1870 | FALSE |
| Santa Margarita River, CA | $\sigma_{obs}$ | 0.587 | 0.478 | 0.706 | 1.001 | 1382 | FALSE |

Table S6: Goodness of fit metrics for the S-TS & LB-TS out-of-sample model predictions for each site.

| | **RMSE ($g\ O_2\ m^{-2}\ d^{-1}$)** | | **NRMSE** | | **Coverage (%)** | |
|---|---|---|---|---|---|---|
| **Site Name** | **S-TS** | **LB-TS** | **S-TS** | **LB-TS** | **S-TS** | **LB-TS** |
| S. Br. Potomac River, WV | 3.34 | 3.43 | 0.22 | 0.23 | 64 | 70 |
| Paint Branch, MD | 1.12 | 0.75 | 0.23 | 0.15 | 91 | 89 |
| Proctor Creek, GA | 1.19 | 1.11 | 0.19 | 0.17 | 76 | 94 |
| Beaty Creek, OK | 1.12 | 1.02 | 0.31 | 0.29 | 86 | 94 |
| Pecos River, TX | 4.16 | 3.51 | 0.27 | 0.23 | 97 | 89 |
| Santa Margarita River, CA | 2.15 | 1.84 | 0.23 | 0.20 | 80 | 85 |
| **Mean** | 2.18 | 1.95 | 0.24 | 0.21 | 82 | 87 |
| **Minimum** | 1.12 | 0.75 | 0.19 | 0.15 | 64 | 70 |
| **Maximum** | 4.16 | 3.51 | 0.31 | 0.29 | 97 | 94 |
